# Standard Intein Gene Expression Ramps (SIGER) for protein-independent expression control

**DOI:** 10.1101/2021.12.07.471673

**Authors:** Maxime Fages-Lartaud, Yasmin Mueller, Florence Elie, Gaston Coutarde, Martin Frank Hohmann-Marriott

## Abstract

Coordination of multi-gene expression is one of the key challenges of metabolic engineering for the development of cell factories. Constraints on translation initiation and early ribosome kinetics of mRNA are imposed by features of the 5’UTR in combination with the start of the gene, referred to as the “gene ramp”, such as rare codons and mRNA secondary structures. These features strongly influence translation yield and protein quality by regulating ribosome distribution on mRNA strands. The utilization of genetic expression sequences, such as promoters and 5’UTRs in combination with different target genes leads to a wide variety of gene ramp compositions with irregular translation rates leading to unpredictable levels of protein yield and quality. Here, we present the Standard Intein Gene Expression Ramps (SIGER) system for controlling protein expression. The SIGER system makes use of inteins to decouple the translation initiation features from the gene of a target protein. We generated sequence-specific gene expression sequences for two inteins (DnaB and DnaX) that display defined levels of protein expression. Additionally, we used inteins that possess the ability to release the C-terminal fusion protein *in vivo* to avoid impairment of protein functionality by the fused intein. Overall, our results show that SIGER systems are unique tools to mitigate the undesirable effects of gene ramp variation and to control the relative ratios of enzymes involved in molecular pathways. As a proof of concept of the potential of the system, we also used a SIGER system to express two difficult-to-produce proteins, GumM and CBM73.

**Graphical abstract:** 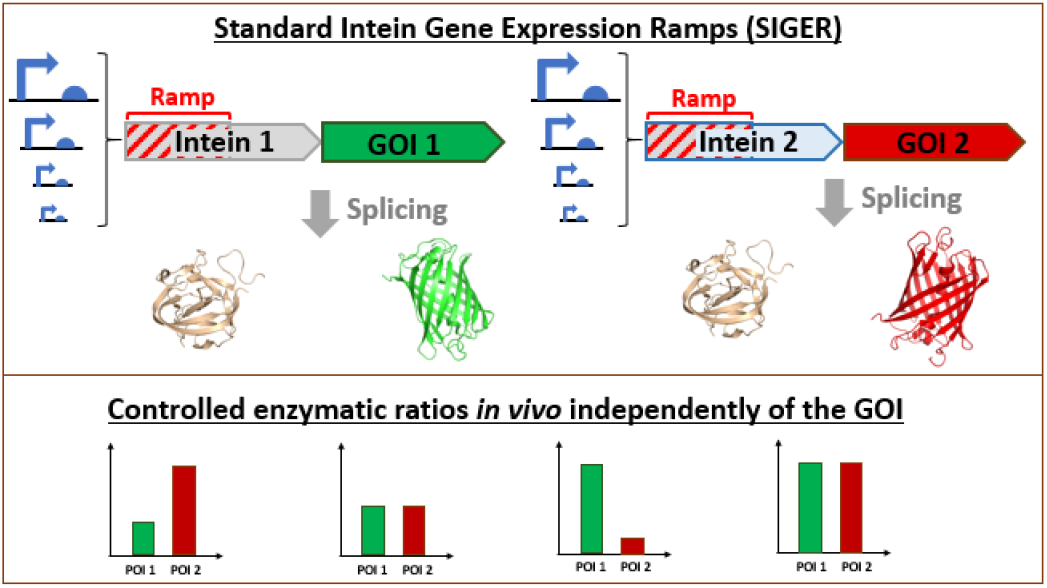

## Introduction

Cell factories are central components of biotechnology for the production of recombinant proteins and biochemicals that find numerous applications in pharmaceutical, agricultural, food, cosmetic and chemical industries^1–3^. The choice of the cellular host depends on the application and is crucial to obtain high product yield^4^ and desired protein properties such as solubility, secretion, glycosylation and other post-translational modifications^5^. The most common cell factories include various bacteria^6, 7^, yeast^8, 9^, filamentous fungi, plants^10^, as well as insect and mammalian cells^11, 12^. The ability of cell factories to produce quality protein in high yields is determined by the type of genetic expression system, the characteristics of the strain and its adaptability to large-scale cultivation processes^13^. For a given host, the choice of genetic expression system is fundamental because it determines the maximum yield, affects protein quality, and protein expression characteristics. For the production of biomolecules, the expression of enzymes involved in a metabolic pathway must be tuned to balance the relative enzymatic activities, thus avoiding burdensome overexpression of proteins and toxicity of intermediate metabolites^14–18^. Unsuitable genetic expression systems can be the source of substantial cell toxicity and protein misfolding leading to the failure of biotechnological endeavors.

Current gene expression systems comprise constitutive and inducible promoters^6, 19^ coupled with native or computationally designed 5’UTRs^20–22^. Single genes are expressed with a monocistronic cassette and multiple genes are expressed with series of monocistronic or polycistronic constructs. For example, the monocistronic pET system (DE3/T7), which is one of the predominant protein expression systems in *E. coli*^23^, consists of an IPTG-inducible T7 polymerase gene integrated into the *E. coli* genome and a pET vector containing a T7 promoter expressing the gene of interest (GOI). However, genetic expression systems are not always successful in expressing certain GOIs or ensuring the functionality of the protein of interest (POI)^6^. The main causes of failure are related to defects in translation, protein folding, protein translocation, mRNA stability, plasmid sustainability and cell viability^6^. The aforementioned transcriptional, translational and protein maturation problems often originate from the genetic expression system and the lack of complementarity between genetic elements or adaptation to the host organism.

Mechanisms governing translation initiation involve mRNA stability, mRNA unfolding and ribosome entry-sites accessibility^24–27^ (*e.g*. Shine-Dalgarno^28^ (SD) or Kozak^29^ sequence). Translation efficiency is highly dependent on the mRNA secondary structures formed within the 5’UTR and between the 5’UTR and the start of the coding sequence (CDS), region referred to as the “gene ramp”, composed of the first 100 to 150 nucleotides^30, 31^, influencing translation initiation and protein yields. The gene ramp influences ribosomal entry onto the mRNA and modulates early translation rates^26,30–36^ (*Figure 1a*). The limitation in early translation rates at 5’ genes termini was hypothesized to allow ribosomal spacing, therefore avoiding traffic jams, ribosome fall-offs and aborted translation^37–39^. The nucleotide composition of the ramp modulates the formation of mRNA secondary structures, in combination with the 5’UTR and within the ramp, that influence translation initiation kinetics^40^. Additionally, the gene ramp is enriched in rare codon clusters that decrease early translation rates^30,31,41,42^ and are involved in determining local mRNA secondary structures^30, 43^. Positively charged amino acids, such as lysine, which are also overrepresented in ramp codons, interact with negatively charged residues in the ribosomal exit tunnel decreasing translation rates^44, 45^. These characteristics of 5’ gene termini properties are linked to conserved evolutionary mechanisms that are shared across many microorganisms^37, 42^.

**Figure 1.**
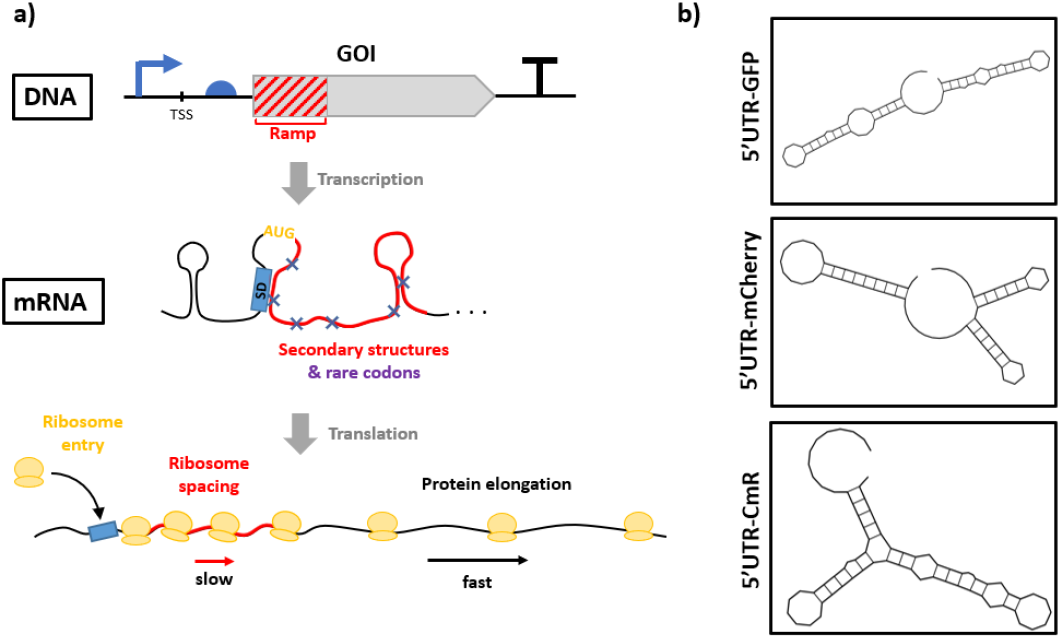
Characteristics of a gene ramp and implications for protein expression. a) Representation of a genetic cassette. The gene ramp corresponds to the first 100 to 150 nucleotides of the gene downstream a 5’UTR. The gene ramp forms secondary structures with the 5’UTR and within itself that imposes constraints on translation initiation. Additionally, rare codons with slow ribosome decoding are over-represented in the gene ramp. These characteristics affect translation initiation and slow early translation rates. The precise control of ribosome kinetics during translation initiation influences translation yield and protein quality. b) Example of the variation in secondary structures formed between an identical 5’UTR and three different coding sequences (±40 bp around the start codon), for *sfGFP*, *mCherry* and *cat*, predicted using the web-based RNAfold tool (http://rna.tbi.univie.ac.at/). The exchange of GOI downstream the 5’UTRs dramatically affects the resulting mRNA secondary structure. The mRNA secondary structures influence the efficiency of translation initiation and protein yield.

The complementarity between the 5’UTR and the gene ramp influences translation yield and protein folding^46–48^. For an established genetic expression system, replacing the GOI changes the resulting secondary structure occurring between the 5’UTR and the ramp (*Figure 1b*). These structural variations affect the outcome of protein production. There are computational tools to help predict 5’UTRs sequences suitable for a given coding sequence^20–22^. However, these tools only provide predictions for individual cases, which have to be experimentally verified, whereas standardized *a priori* experimentally-validated systems would be preferable. In addition, codon optimization of GOIs can affect the composition of rare codons within the ramp^42^, leading to perturbations in translation rates that produce insoluble or misfolded proteins^46–48^.

The principal solution deployed to circumvent incompatibility between genetic elements and rescue protein solubility is the use of N-terminal fusion protein tags. These tags increase the POI’s solubility, provide a compatible buffer sequence with genetic expression parts, and some tags can be used for downstream purification process^49, 50^. Most solubility tags require the use of specific proteases to liberate the protein of interest (POI) from the fusion construct^51^. One exception is the family of 2A self-cleaving linker peptides^52^ that display the ability to excise themselves from a fusion protein, releasing the POI from the fusion tag. Inteins are another type of self-cleaving proteins that are used in protein purification^53, 54^ (*e.g*. the IMPACT system from New England Biolabs). Inteins are naturally occurring autocatalytic proteins that possess the ability to excise themselves from a larger protein, ligating the two flanking proteins (exteins) together in the splicing process^55^. The protein splicing process is spontaneous, occurs post-translationally and does not require the intervention of exogenous factors or proteases^55^. In the case of the IMPACT system, the *Sce* VMA intein is combined with a chitin-binding domain to bind the appropriate resin, and exhibit N-terminal cleavage in the presence of DTT or β-Mercaptoethanol^56^. Split inteins are another type of inteins, composed of an N-intein and C-intein, each fused to an extein, individually translated. After, translation, the N- and C-intein fragments assemble non-covalently to provide the canonical intein structure and carry out *trans* protein splicing^55, 57^. In addition, inteins have been engineered to selectively release the N- or C-terminal peptide by mutating key catalytic residues. These unique properties enable a wide variety of applications such as enzyme activation, protein ligation, production of cyclic peptides, protein purification systems, biosensors and reporter systems^55,58–60^. One example of such an application is the simultaneous production of two equimolar POIs by using a dual-intein system composed of the respective N- and C-terminal cleavage properties of *Ssp DnaE* and *Ssp DnaB*^61^.

Here, we used C-terminal cleaving inteins to design Standard Intein Gene Expression Ramps (SIGERs). Mini-inteins are short genes that can provide the necessary properties of a gene ramp while releasing the POI *in vivo*. We used the geneEE method^62^ to create artificial promoter/5’UTRs that interact with the intein sequences in order to obtain a wide variety of gene ramps displaying a broad range of protein expression levels. This way, our SIGER systems fulfill the genetic complementarity requirements of a gene ramp, thus ameliorating protein folding and solubility, as well as offering the possibility to adapt expression levels to avoid cellular burden. Since translation initiation imposes major constraints on protein yield and protein quality, SIGER systems offer a buffer genetic region that allows exchanging GOIs without affecting the level of protein expression. Furthermore, we show that coupling different SIGER systems in the expression host permits facile control of enzymatic ratios to balance metabolic pathways. The standard intein systems tightly controls the production of discrete enzymes *in vivo* in the desired quantity, without concerns for complementarity between genetic expression elements. Overall, we developed two SIGER systems and explored their ability to fulfill ramp properties and control multi-enzyme expression levels.

## Results and Discussion

### Design of the standard intein gene expression ramp (SIGER)

The principle of a standard gene expression ramp is to buffer the complementarity effect of the gene expression sequences (GES), composed of a promoter + 5’UTR, with the gene of interest. We selected two C-terminal cleaving mini-inteins, *DnaB*^63^ and *DnaX*^64^, of 159 and 140 amino acids, respectively, to serve as gene expression ramps (for sequences see *Supplementary Table S2*). The length of the intein genes is around 450 bp, which is long enough to absorb the properties of a ramp that affects usually the first 150 bp after the start codon (*Figure 2a*). The POI is released from the intein by autocatalytic cyclization of the C-terminal Asn residue (*Figure 2b*), N159 and N140 for DnaB and DnaX respectively. The efficiency of the reaction is dependent on the residue adjacent to the C-terminal asparagine. Ser-Arg residues at the C-terminal end of inteins have been experimentally demonstrated to result in high C-terminal self-cleavage activity^65^. Therefore, a C-extein amino acid linker, SRGP^61^, was added to the C-terminal end of DnaB and DnaX. In this way, the intein gene fragment should function as the *in vivo* gene ramp and the resulting intein translation product can release the C-terminal fusion protein of interest (POI). Using SIGER, a discrete POI is produced, and thus avoiding potential defects in functionality due to the steric hindrance by the N-terminal fusion protein.

**Figure 2.**
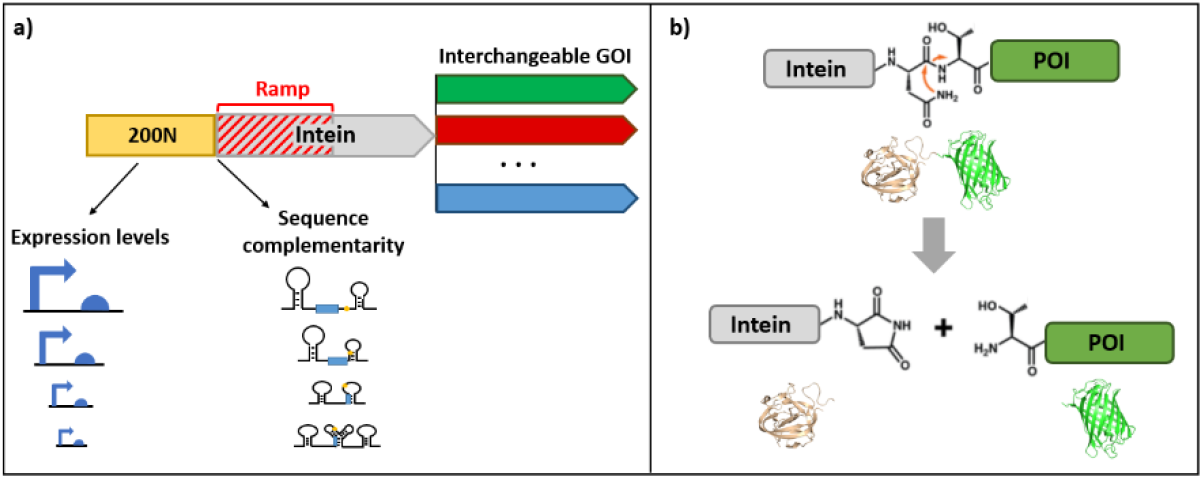
Characteristics of the standard intein gene expression ramp (SIGER) systems. a) Genetic organization of SIGER systems. The inteins fulfill the role of standard gene ramps and the 200N random DNA fragment provides intein-tailored gene expression sequences (GES). The nucleotide sequence of GES 5’UTRs produces different mRNA secondary structures in relation to the sequences of the inteins and each GES displays various expression levels. b) Autocatalytic C-terminal cleavage of inteins. The cyclization of the N-terminal Asn residue releases the POI *in vivo*.

We first aimed to create artificial GES tailored to the *DnaB* and *DnaX* sequences, which possess a wide range of expression. For this, we used the gene expression engineering method (GeneEE) that consists in placing a DNA segment of 200 random nucleotide (200N) directly upstream a protein-coding sequence^62^. GeneEE provides a functional GES for 30 to 40% of sequences, a wide range of expression and gene-tailored 5’UTRs in *E. coli*^62^. The application of GeneEE created a multitude of GES tailored to the *DnaB* and *DnaX* sequences that displayed different levels of expression (*Figure 2a*).

The combined properties of inteins and the GeneEE method can provide interesting gene expression tools that define expression levels independently of the GOI sequence. The standard intein gene expression ramps (SIGER) circumvent issues related to the complementarity of gene ramps and GOI, as well as release discrete proteins. The GeneEE method can provide intein-tailored GES candidates with fine-tuned protein expression characteristics that limit cell toxicity due to burdensome protein expression. Finally, the combination of characterized SIGER systems can permit to define the level of expression of different POIs to balance metabolic pathways.

### Fabrication of intein-tailored promoter library

We used the GeneEE method to generate a library of GES adapted to the sequence of *DnaB* and *DnaX*. To do so, we inserted the 200N DNA fragment in front of *DnaB-GFP* and *DnaX-GFP* by Golden Gate assembly and transformed it into *E. coli* (see *Methods* and *Supplementary Figure S1*). Positive clones (n=186) displaying seemingly green fluorescence under UV light were selected and grown overnight in 96 well plates. The fluorescence measurement of the 186 positive clones for each *DnaB-GFP* and *DnaX-GFP* are presented in *Figure 3*. The artificial GES resulted in a wide range of expression levels spanning over one order of magnitude. For each intein, 10 strains were selected to represent characteristic levels of expression. After eliminating strains showing inconsistencies, such as strains with large variability in GFP fluorescence levels, we obtained seven and eight gene-tailored GES for *DnaB* and *DnaX*, respectively (GES were named B_P1 to B_P7 for *DnaB* and X_P1 to X_P8 for *DnaX*) (*Figure 4*). All the GES identified with *DnaB-GFP* and *DnaX-GFP* were sequenced (*Supplementary Table S3*). Using the complimentary GES and intein ramps, we investigated the effect of exchanging the GOI on the gene expression levels.

**Figure 3.**
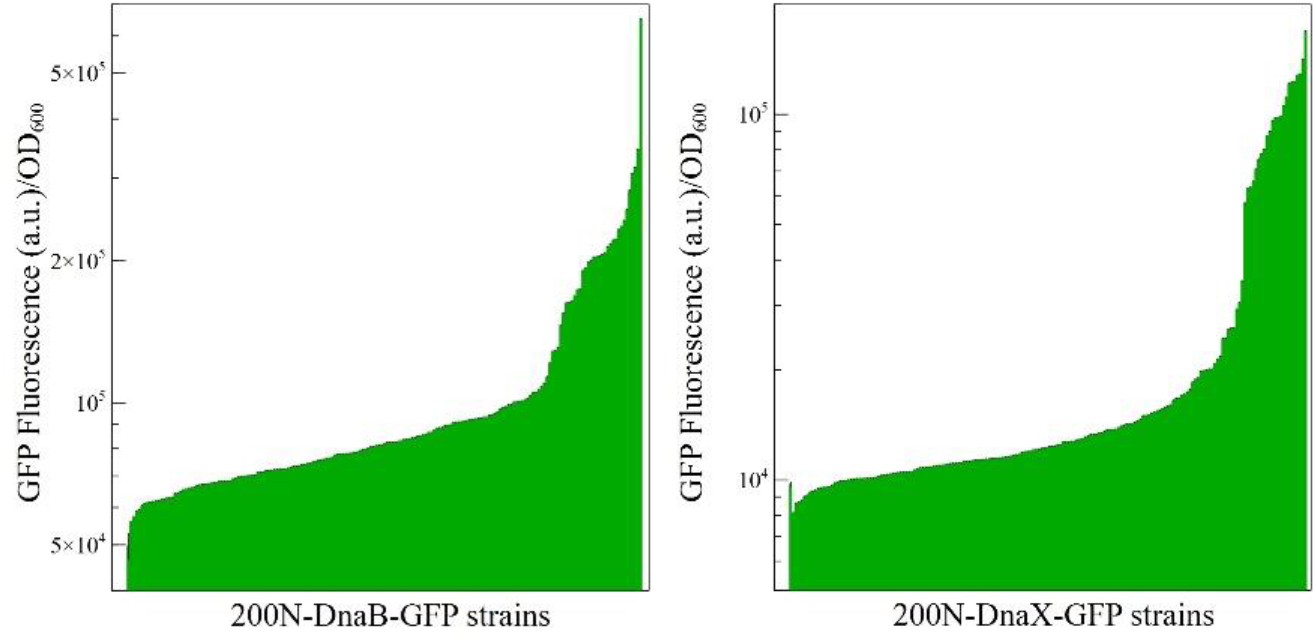
Fluorescence intensity of the GES library tailored for *DnaB* and *DnaX*. The histograms represent single fluorescence measurements for the GES library expressing *DnaB-GFP* and *DnaX-GFP*. For each intein, 186 strain expressing various levels of GFP were analyzed. The first histogram bar (position 1) represents the average auto-fluorescence of a negative control performed in triplicates and standard deviation is shown in black (calculated for this sample only).

**Figure 4.**
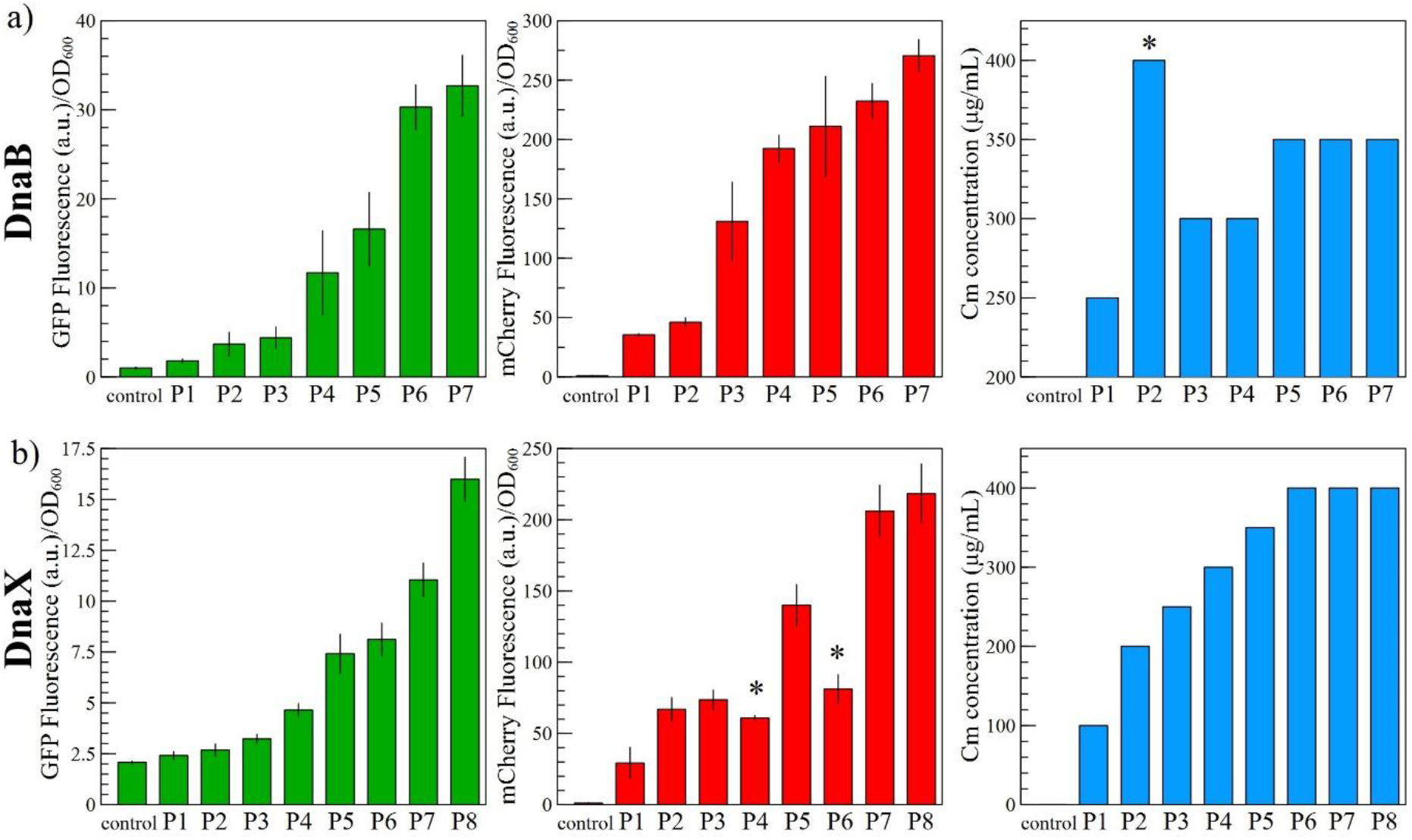
Expression of different GOIs with the DnaB and DnaX SIGER systems. The GES specific for *DnaB* and *DnaX* were transferred to two other gene cassettes to express *mCherry* and *cat*. The fluorescence measurements of GFP (green) and mCherry (red), and the chloramphenicol resistance profile of each strain (blue) are presented for the a) *DnaB* and b) *DnaX* SIGER systems. The expression range selected with GFP is generally conserved when a different GOI is placed after the inteins. The reasons for the observed exceptions (indicated with an asterisk) are discussed in the main text.

### Gene expression intensity is independent of the GOI

Ten strains for each inteins were selected to represent distinct thresholds values of GFP protein expression (*Figure 3*). From these 10 strains, the levels of gene expression for seven (*DnaB*) and eight (*DnaX*) strains were defined as conserved because they showed consistent fluorescence levels when the measurements were performed in triplicates (*Figure 4*). In order to evaluate the genetic buffering effect of SIGER systems, we expressed *mCherry* and the chloramphenicol resistance gene *cat*. The GOIs *mCherry* and *cat* were inserted downstream both inteins (see *Methods* and *Supplementary Figure S1*), to create *DnaB-mCherry, DnaB-cat, DnaX-mCherry* and *DnaX-cat*. Then, we extracted the seven and eight GES identified with *DnaB-GFP* and *DnaX-GFP* respectively to and placed them in front of the new gene cassettes (see *Methods* and *Supplementary Figure S1*). Each construct was transformed into *E. coli* and positive clones were grown overnight in triplicates in 96 well plates. The fluorescence of clones containing *DnaB-mCherry* and *DnaX-mCherry* was quantified (*Figure 4*). The strains containing *DnaB-cat* and *DnaX-cat* were diluted 100 times and replica-plated on LB-agar plates supplemented with increasing chloramphenicol concentrations to evaluate their resistance profile (*Figure 4* and *Supplementary Figure S2*). The fluorescence of strains carrying weak GES was confirmed by fluorescence microscopy (*Supplementary Figure S3*).

For *DnaB-mCherry*, a first observation is that fluorescence levels are generally higher than expected from the *DnaB-GFP* range although the relative strength ranking of GES remained the same (*Figure 4a*). This slight *mCherry* overexpression could be due to the newly identified alternative translation start site of *mCherry* that produces a short functional protein isoform^66^. The short mCherry isoform produces significant background fluorescence when the reporter is used as a C-terminal fusion partner. Although the background fluorescence is supposedly the same across strains, the production of the short mCherry isoform blurs the relative differences in the expression range.

The chloramphenicol gradient also confirms the relative strength of each *DnaB*-specific GES (*Figure 4a*), with the exception of B_P2 that presents a high chloramphenicol resistance (400 μg/mL) when the expectations would place it closer to B_P1 (250 μg/mL) in respect to fluorescence measurements of GFP and mCherry. Sanger sequencing of B_P2 from *DnaB-cat* revealed that the GES was a different sequence than the GES in B_P2 from *DnaB-GFP* (*Supplementary Table S3*). This discrepancy can be explained in the following way, when using the GeneEE method, after cloning of the 200N DNA fragment, several plasmids can be transformed into one *E. coli* cell. The different GES can be sustained in one *E. coli* cell due to the relatively small nucleotide variation of the plasmid (only in the 200N region). After prolonged growth, different cell populations can emerge bearing only one of the plasmid isoforms. Here, the PCR amplification of the GES regions of *DnaB-GFP* may have extracted different GES copies. The subsequent cloning onto new gene cassettes can yield strains with an unexpected sequence. This effect yielded two different *DnaB-P2-cat* strains with different representation in the population. The selection on chloramphenicol favored the underrepresented strain that possessed a stronger promoter.

For *DnaX-mCherry*, the expression range is conserved with respect to *DnaX-GFP* except for X_P4 and X_P6 (*Figure 4b*). As for *DnaB-P2-cat*, Sanger sequencing of *DnaX-P4-mCherry* revealed a different DNA sequence than *DnaX-P4-GFP*, thus explaining the lower fluorescence than expected. However, for *DnaX-P6-mCherry*, DNA sequencing provided the same sequence as *DnaX-P4-GFP*. The fluorescence defect of *DnaX-P6-mCherry* may originate from another random mutation on the plasmid. The chloramphenicol resistance profile of *DnaX-cat* strains confirms the expression range obtained with *DnaX-GFP*, although it was not possible to differentiate between expression levels for X_P6, X_P7 and X_P8 because of insufficient method sensitivity (*Figure 4b*).

Overall, the expression range of the SIGER systems established with *GFP* were conserved when the GOI was exchanged with *mCherry* or *cat*. Apart from the sustained multiple GES issue, data shows that the different GES coupled to each intein provide standard gene expression cassettes with defined, consistent and predictable translation levels. We then investigated the combination of SIGER systems to tune multi-enzyme expression.

### Coupling SIGER systems for balancing multi-gene expression in vivo

The developed SIGER systems are unique tools that enable fine-tuning of the expression of different discrete enzymes in the desired quantity *in vivo*. In order to test the accuracy of SIGER systems in controlling multi-enzyme expression, we coupled the *DnaB-mCherry* and *DnaX-GFP* cassettes on the same plasmid. The two intein cassettes were coupled on a level 2 vector (Lv2), containing a chloramphenicol resistance marker, by using the level assembly method developed by Fages-Lartaud *et al*. for pathway assembly^67^ (see *Methods* and *Supplementary Figure S1*). In brief, the *DnaB* and *DnaX* gene cassettes were constructed on level 1 plasmid (Lv1), bearing an ampicillin marker, and containing BbsI restriction sites on each side of the gene cassettes. Then, the complementarity of the BbsI scars was used to assemble each cassette on the Lv2 vector. As a demonstration, we selected three GES for each intein that represent distinct levels of expression. For *DnaB-mCherry*, we used B_P2 as a weak GES, B_P3 as an intermediate GES and B_P7 as a strong GES. For *DnaX-GFP*, we used X_P2 as a weak GES, X_P5 as an intermediate GES and X_P7 as a strong GES. Each *DnaB* cassette was coupled to the three *DnaX* cassettes, resulting in nine genetic combinations. Each coupled construct was transformed into *E. coli* and cells were selected on LB-agar plates supplemented with 30 μg/mL chloramphenicol. The *E. coli* strains carrying each of the nine constructs were grown overnight in triplicates in 96 well plates and the fluorescence of mCherry and GFP was measured for each strain. The results are presented in *Figure 5*.

**Figure 5.**
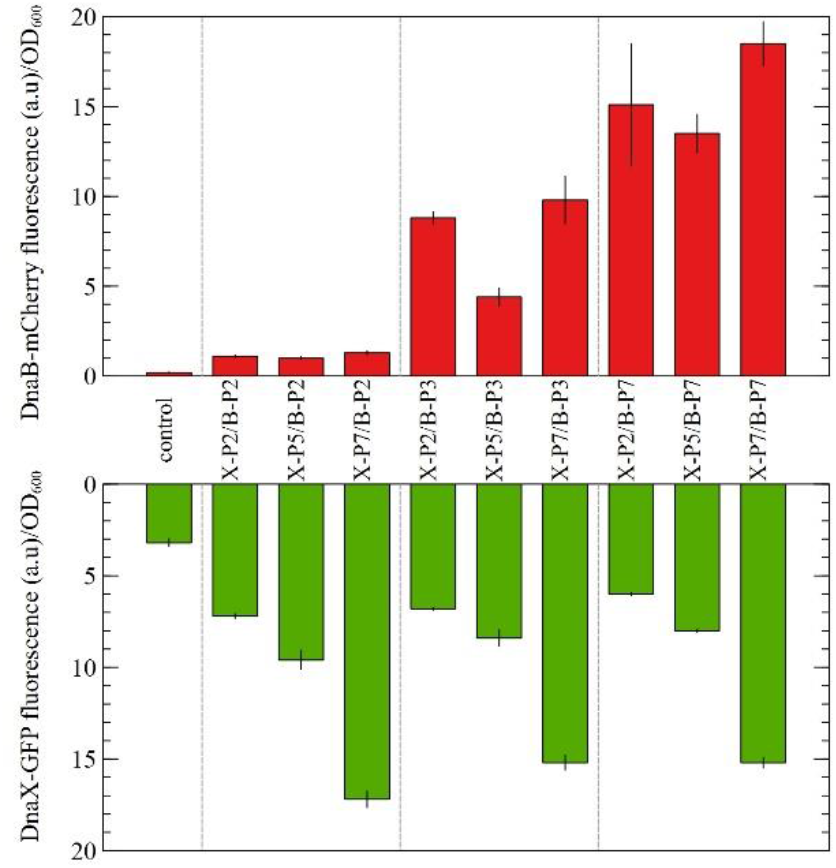
Controlled enzymatic expression by coupling two SIGER systems. The *DnaB-mCherry* and *DnaX-GFP* gene cassettes, each expressing at three different levels, were coupled on the same plasmid to assess the ability of SIGER systems to control multi-enzymatic expression. The histograms represent the fluorescence levels of the weak, intermediate and strong GES (respectively B_P2, B_P3 and B_P7 for DnaB; and X_P2, X_P5 and X_P7 for DnaX). The SIGER systems present very good control over the protein expression of any POI *in vivo*.

The first observation is that the expression intensities of *DnaB-mCherry* and *DnaX-GFP* were conserved when compared to single cassette measurements (see *Figure 4*), with the exception of B_P3 in the coupled system possessing B_P3 and X_P5. DNA sequencing revealed that B_P3 of that specific construct contained an unexpected sequence, thus explaining the discrepancy in fluorescence intensity. Beside this issue, the *DnaB* SIGER displayed three distinct levels of expression from low expression to 8.2 and 13.8-fold increase (*Figure 5)*. The *DnaX* SIGER also showed three expression levels with a 1.6 and 3.7-fold increase relatively to the weak GES (*Figure 5)*. The fluorescence of weak GES and the cellular co-localization of *mCherry* and *GFP* was confirmed by fluorescence microscopy (*Supplementary Figure S3*).

Secondly, results show that coupling SIGER systems enables control of the level of expression of two discrete proteins. The nine GES combinations of the *DnaB* and *DnaX* SIGER systems present various levels of relative enzymatic expression. Based on these results, we show that conceivably SIGER systems may be applied to control the expression of enzymes in a metabolic pathway to limit cellular burden or toxicity without the need to engineer specific GES for each GOI.

The last verification regarding the functionality of SIGER systems was to confirm the ability of the C-terminal cleaving inteins to release the POI. Indeed, one of the objectives of the SIGER systems is to produce *in vivo* discrete POIs without interference from an N-terminal fusion partner. Therefore, we investigated the efficacy of C-terminal cleavage by *DnaB* and *DnaX in vivo*.

### Assessment of in vivo cleavage

In order to assess the efficiency of intein-mediated *in vivo* C-terminal cleavage, we created two mutants of *DnaB* and *DnaX* possessing an N-terminal truncation (see *Methods*). The truncation renders the intein cleavage non-functional by preventing the N-terminal asparagine cyclization resulting in the release of the POI. *DnaB-GFP* and *DnaX-GFP*, expressed by a strong GES, and their mutated versions were transferred into *E. coli* BL21 for protein production at 30°C in canonical Erlenmeyer flasks. Cell cultures were harvested by centrifugation, cell content was released by sonication and cell debris were eliminated by centrifugation. Proteins present in the soluble fractions were identified by SDS-PAGE followed by Western Blot to detect the 6xHis tag of GFP. Furthermore, soluble fractions were chromatographically purified using a HisTrap HP Ni-sepharose column and evaluated on SDS-PAGE prior to LC-MS analysis (*Supplementary Figure S4)*.

The Western Blot results are presented in *Figure 6*. For *DnaB-GFP*, a clear GFP band is observable at 28 kDa and a faint band is noticeable around 45 kDa. This result shows that DnaB in vivo cleavage is efficient and that the *DnaB*-SIGER system releases the POI. The 45-kDa protein corresponding to the uncleaved DnaB-GFP fusion protein was analyzed by LC-MS after protein purification and SDS-PAGE. The analysis confirms that the 45-kDa protein is the uncleaved DnaB-GFP fusion protein (see *Supplementary Figure S5*).

**Figure 6.**
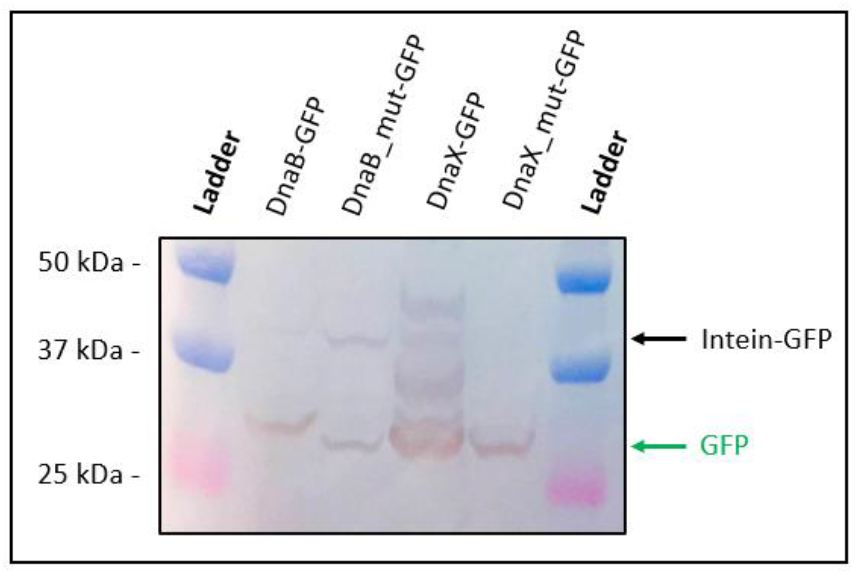
Western blot of the soluble protein fractions of the *DnaB* and *DnaX* SIGER systems and their truncated versions. The molecular weight of the bands in the ladder are indicated on the left side and sample names are indicated on top. The uncleaved Intein-GFP protein and the released free GFP are indicated with black and green arrows respectively. The DnaB-GFP protein displays efficient dissociation. The truncated version, DnaB_mut-GFP, shows the expected size of an uncleaved fusion protein. The DnaX-SIGER system presents several bands including uncleaved fusion protein, free GFP and intermediate bands. The origin of these intermediate bands is discussed in the text, but they were absent from the SDS-PAGE (*Supplementary Figure S4*). Regardless, the DnaX-GFP protein shows only partial *in vivo* cleavage. The GFP bands present for the mutated DnaB and DnaX versions originates from alternative translation start sites that were confirmed by LC-MS analysis.

The mutated version of *DnaB-GFP* displays a clear 43-kDa band corresponding to the uncleaved fusion protein, but also an unexpected 28-kDa protein around the size of GFP. This last-mentioned protein might be the result of an alternative translation start site occurring just upstream the *GFP* sequence. The mutated DnaB-GFP protein does not contain internal methionine residues that could result in the production of a GFP isoform. However, there are a several valine residues, such as V136 and V139, that could serve as alternative start codons, especially given the presence of an upstream SD sequence (GGAG), six nucleotides upstream of V136. LC-MS analysis of the purified unexpected 28-kDa protein confirmed that it is GFP with a small N-terminal peptide elongation (at least DLTVPGPR) (see *Supplementary Figure S5*). Alternative translative start sites are a common issue occuring with N-terminal protein tags and fluorescent proteins^66, 68^. This result support the hypothesis of an alternative start codon close to the gene start producing a GFP isoform. Nevertheless, this artifact does not interfere with the conclusion regarding the *in vivo* cleavage performance of DnaB-GFP.

For *DnaX-GFP*, immunoblotting presents several bands, a strong 28-kDa band of the discrete GFP protein, a lighter 43-kDa band related to the uncleaved DnaX-GFP protein, and a couple intermediate bands (*Figure 6*). The relatively strong intermediate band may originate from an internal alternative translation start site created by the methionine residue M77 of DnaX that results in a 35-kDa protein. The intermediate isoforms were not present on the SDS-PAGE performed after purification (*Supplementary Figure S4*). The Western Blot and the SDS-PAGE performed with purified fractions both suggest an incomplete cleavage of the DnaX-GFP fusion protein. However, the previous authors that engineered DnaX did not notice such defect in protein cleavage, which could be resulting from our experimental setup. The reason for the low cleavage efficiency may be either low cleavage kinetics or suboptimal protein production conditions for full cleavage of DnaX-GFP. The 43-kDa protein was confirmed to correspond to the uncleaved DnaX-GFP fusion protein by LC-MS analysis (see *Supplementary Figure S5*).

The truncation of *DnaX* introduces a stop codon at the start of the *GFP* gene by creating a frameshift. Unfortunately, an in-frame GFP isoform can still be produced from the mutated *DnaX-GFP* mRNA due to the presence of an unexpected methionine located nine amino acids before the start of the *GFP* gene. In addition, an SD-like sequence, GGAA, is present seven nucleotide before the mentioned methionine. The combination of these elements resulted in the production of a GFP isoform possessing a nine amino acid N-terminal extension (*Figure 6*).

The analysis of intein C-terminal cleavage revealed a relatively good *in vivo* cleavage of DnaB and GFP and an incomplete excision of DnaX from the fusion construct. Intein autocatalytic cleavage is highly dependent on conditions such as maturation time, pH, salinity, chemical additives and temperature^57,69–71^, depending on the origin of the intein. Prior to protein purification, it is possible to adjust the buffer composition to facilitate the release of the POI. DTT is commonly added in intein-based purification systems. A decrease in pH from 7.5 to 6.0 significantly increase intein C-terminal cleavage^69^. However, in physiological conditions, temperature is the principal parameter that can be modified without impairing cell growth. It is important to note that full cleavage is not necessarily a condition to yield a functionaly active POI.

Therefore, we investigated the effect of temperature on protein production and intein cleavage. An engineered strain containing DnaB-GFP expressed by a strong constitutive promoter (P7) was cultivated at 22°C and 37°C. The His-tag-containing GFP was purified on a Nickel-sepharose column and analyzed on SDS-PAGE (*see **Figure 7** and Supplemental Table S4*). The first outcome was that a higher temperature was correlated with higher protein yield, 32 mg/L at 37°C and 5 mg/L at 22°C. Secondly, we estimated that intein cleavage was more efficient at 37°C (64%) than at 22°C (43%). The Western blot results from *Figure 6* suggest even higher cleavage efficiency, but this could be explained by the maturation time of the fusion protein; indeed, studies showed that cleavage takes one to a few hours to be complete *in vitro*^57^.

**Figure 7.**
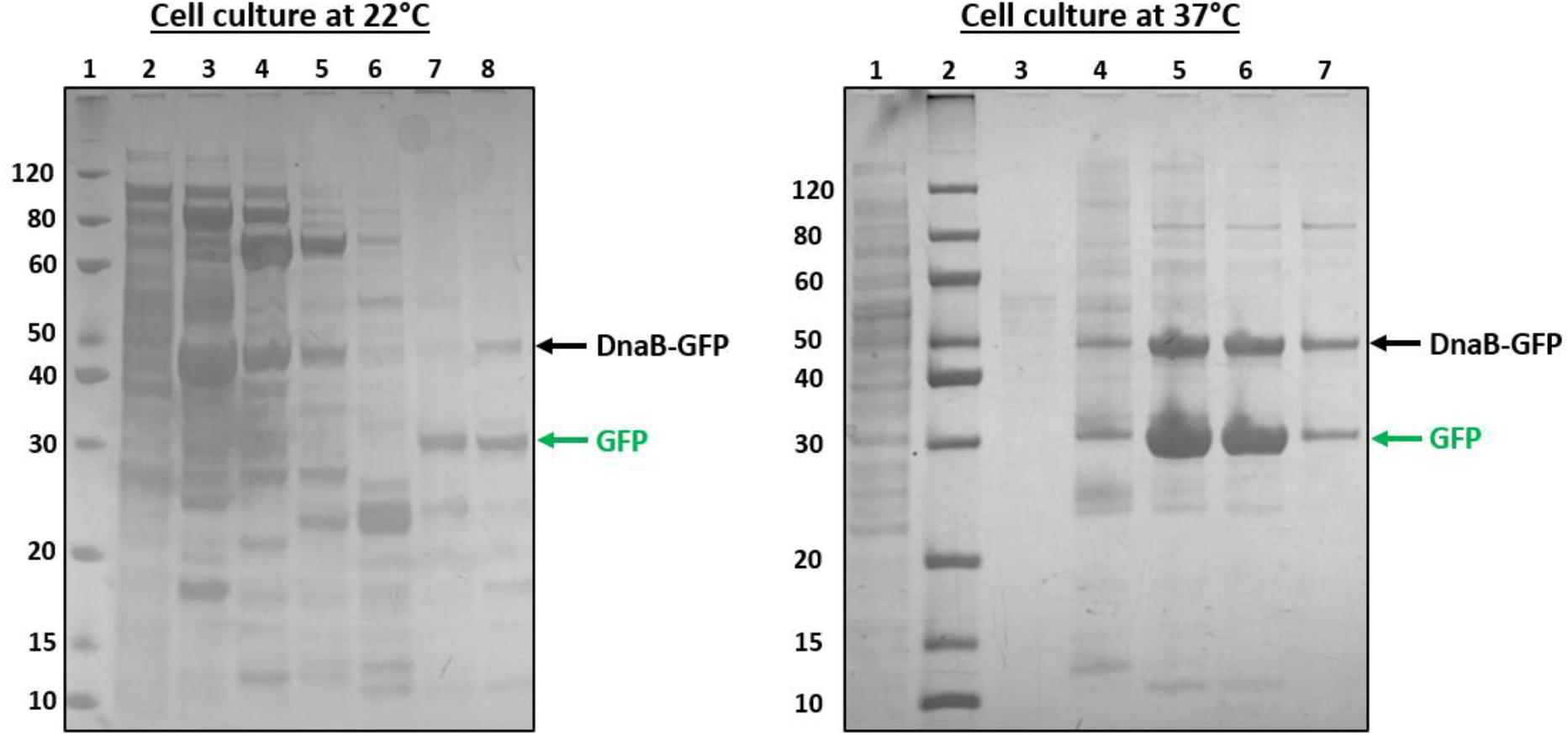
Influence of growth temperature on intein cleavage. SDS-PAGE showing Nickel purified fractions of DnaB-GFP produced in E. coli cultivated at 22°C (left) and 37°C (right). Lanes of 22°C (left) gel: 1: ladder, 2: soluble fraction, 3 to 8: Ni-column elution fractions. Lanes of 37°C (right) gel: 1: soluble fraction, 2: ladder, 3 to 7: Ni-column elution fractions. The uncleaved DnaB-GFP protein is indicated with a black arrow, and the cleaved GFP is indicated with a green arrow. Intein cleavage efficiency was estimated with ImageJ at 64% for the 37°C culture and at 43% for the 22°C culture. Calculations included cleaved and uncleaved protein from all purified fractions. Protein yields was estimated from the chromatogram at 5 mg/L and 32.1 mg/L for 22°C and 37°C cultures respectively (*Supplemental Table S4*).

**Figure 8.**
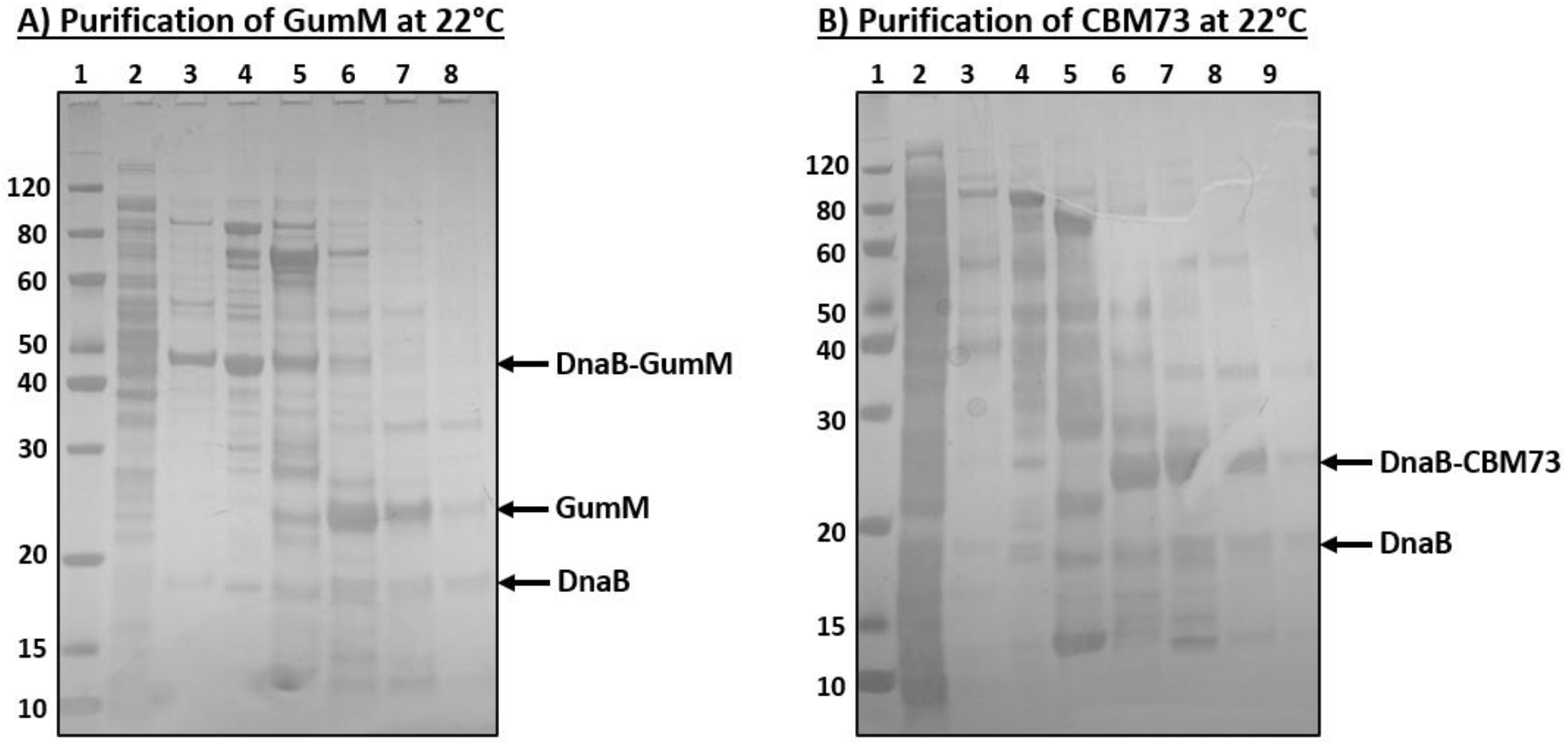
Production and purification of the difficult-to-express proteins GumM and CBM73. SDS-PAGE showing Nickel purified fractions of A) DnaB-GumM and B) DnaB-CBM73 produced in E. coli at 22°C. A) lane 1: ladder, 2: soluble fraction, 3 to 8: Ni-column elution fractions. B) lane 1: ladder, 2: soluble fraction, 3 to 9: Ni-column elution fractions. Intein cleavage efficiency was estimated with ImageJ at 51% for GumM and at 37% for CBM73. Calculations included cleaved and uncleaved protein from all purified fractions. Protein yields was estimated from the chromatogram at 3.6 mg/L and 2.9 mg/L for GumM and CBM73 respectively (*Supplemental Table S4*).

### Expression of difficult-to-produce proteins with SIGER systems

In this section, we show that SIGER systems can facilitate the expression of difficult-to-produce proteins and yield soluble enzymes. To do so, we aimed to produce two proteins, the glycosyltransferase GumM from *Kozakia baliensis*^72^, involved the biosynthesis of xanthan gum; and the carbohydrate-binding module CBM73 of a lytic polysaccharide monooxygenase from *Cellvibrio japonicus*^73^. A previous investigation of GumM did not succeed in producing the protein using a pTYB1 expression vector, which is based on a C-terminal intein tag, for the IMPACT-CN purification system^74^. Previously, CBM73 was produced only in a low yield of 2 mg/L from a pNIC-CH plasmid containing a T7 promoter coupled with a strong RBS^73^. These proteins are considered difficult to produce because GumM is naturally membrane associated and CBM73 is small and has a hydrophobic surface, both these features can lead to post-translational aggregation, which explains the previously observed low yields.

Here, we used the *DnaB* SIGER system under the constitutive P7 promoter and inserted the *GumM* and *CBM73* genes in place of *GFP*. We successfully produced both GumM and CBM73 protein in yields estimated to 3.6 and 2.9 mg/L respectively. The production was performed at 22°C, which also resulted in low yield for GFP (5 mg/L), hence, these results are promising as production can be improved by almost one order of magnitude by adjusting parameters such as growth temperature and maturation time before downstream purification process. The DnaB-SIGER system enabled for the first time the production of GumM, and improved the production of CBM73 to a yield compatible with downstream utilization and characterization. In this experiment, the pH value of the buffer was set at 8.0, it is possible to improve release of the POI by decreasing the pH between 6 and 7.

## Conclusion

In this study, we used the self-cleavage properties of inteins to create gene expression systems that allow standardization of expression for different proteins. With the GeneEE method, we generated libraries of GES tailored to *DnaB* and *DnaX* sequences. A subset of GES was selected to represent distinct expression levels. These GES coupled to *DnaB* and *DnaX* constitute SIGER systems that control protein expression levels independently of the GOI. The GeneEE method provides a promoter and a 5’UTR tailored to the intein sequence; one of its strength is to be applicable across various bacteria. Therefore, once established, SIGER systems may be applied across a wide range of organisms. Although the selected expression range of GES may be limited, we demonstrate that SIGER systems can be used to balance the expression of multiple enzymes in vivo. The main advantage of using inteins in the SIGER system is that the POI is released from the fusion construct *in vivo*, precluding interference by fused protein tags on enzyme activity. Moreover, the intein gene ramp ensures optimal complementarity of genetic expression sequences with GOIs. The N-terminal intein tag may also favor solubility of the downstream POI as we demonstrated for the production of difficult-to-produce proteins. For protein production, SIGER systems could benefit from being couple to an inducible promoter and a synthetic strong RBS. It is also the case for regulating *in-vivo* enzymatic ratio, as 5’UTR/CDS complementarity affect expression levels, SIGERs constitute a buffer sequence that circumvent the effects of mRNA structures. The *DnaB* SIGER system showed efficient *in vivo* cleavage, while the *DnaX* SIGER system presented incomplete cleavage. Inteins cleavage occurs spontaneously and is not host-specific. The release of POI from an intein is dependent on temperature, pH, maturation time, salinity, chemical additives; these parameters can be adapted during protein production or the purification process. The methodology presented here can be applied to other C-terminal cleaving inteins to create new SIGER systems that could be superior to the *DnaB* and *DnaX* SIGER used in this study or allow coupling of several SIGER systems. We present a non-exhaustive list of mini inteins that have the potential to become SIGER systems (***Table 1***). Nature provides a wide variety of mini-inteins with conserved characteristics and some variability^57,60,75^, for example, the amino acid following the C-terminal asparagine is usually a serine, threonine or cysteine, one of which can have a better cleavage efficiency than the other in a particular intein-context^69^. In addition, the N-terminal cysteine of the mini inteins may need to be mutated to alanine to avoid linkage of N-extein residue if present. SIGER systems represent a new synthetic biology tool that will open new doors for applications in the expression of proteins and investigation of metabolic pathways.

**Table 1.** List of mini inteins suitable for design of new SIGER systems.

| Mini intein | Origin | Length<br>(amino acids) | comments | Reference |
| --- | --- | --- | --- | --- |
| *Ssp DnaB* | *Synechocystis* sp.<br>*PCC6803* | 159 aa |  | Zhang et al.<sup>61</sup> |
| *Ssp DnaX* | *Synechocystis* sp.<br>*PCC6803* | 144 aa | Contains internal<br>his tag | Qi et al.<sup>76,77</sup> |
| *Mtu RecA* | *Mycobacterium*<br>*tuberculosis* | 137 aa |  | Hiraga et al.<sup>78</sup> |
| *Npu DnaE* | *Nostoc punctiforme*<br>*PCC 73102* | 110 aa |  | Xia et al.<sup>79</sup> |
| *RadA-min* | *Pyrococcus*<br>*horikoshii* | 168 aa | thermophile | Hiltunen<sup>80</sup>,<br>Oeemig<sup>81</sup> |
| *Ter DnaE-3* | *Trichodesmium*<br>*erythraeum* | 142 aa | Contains internal<br>his tag | Lin et al.<sup>57</sup> |
| *Ter ThyX* | *Trichodesmium*<br>*erythraeum* | 170 aa |  | Lin et al.<sup>57</sup> |
| *CneA Prp8* | *Cryptococcus*<br>*neoformans* | 178 aa |  | Lin et al.<sup>57</sup> |
| *Ssp GyrB* | *Synechocystis* sp.<br>*PCC6803* | 162 aa |  | Lin et al.<sup>57</sup> |
| *Tvo Vma* | *Thermoplasma*<br>*volcanium* | 170 aa | thermophile | Hiltunen<sup>80</sup> |

## Acknowledgments

This study was supported by a PhD fellowship awarded to M.F-L by the Faculty of Natural Sciences of the Norwegian University of Science and Technology. This work was financed by the Novo Nordisk Foundation (Grant NNF18OC0032242). We thank our partners at Vectron Biosolutions AS for technical assistance. We are grateful to Prof. Dr.-Ing. Jochen Schmid for sharing with us the Kb_pSRK plasmid harboring the *K. baliensis* GumM gene.

## Conflict of interest

The authors declare no conflict of interests.

## Material and Methods

### Materials

*Escherichia coli* DH5-α (New England Biolabs) was used as the cloning and testing strain in this work. *Escherichia coli* BL21 was used as the protein expression strain for downstream protein purification. Cells were grown in LB-Lennox (10 g/L casein peptone (Oxoid, ThermoFisher Scientific), 5 g/L yeast extract (Oxoid, ThermoFisher Scientific), 5 g/L NaCl (VWR) and supplemented with 15 g/L agar (Oxoid, ThermoFisher Scientific) for agar plates) supplemented with the appropriate antibiotics (Sigma-Aldrich). All enzymes were purchased from New England Biolabs. Primers were ordered from Eurofins Genomics or Sigma-Aldrich (primer list: *Supplementary Table S1*). PCR reactions were performed with Q5 polymerase (NEB) unless specified otherwise. Colony PCR reactions were performed with *Taq* polymerase (NEB). QiaQuick PCR purification kit (Qiagen) and QIAprep plasmid Miniprep kit (Qiagen) were used for the purification of PCR products and plasmid DNA respectively. DNA sequences were confirmed by Sanger sequencing performed by Eurofins Genomics.

### Construction of standard cassettes

Genes encoding the mini-inteins ssp-DnaB^63^ and ssp-DnaX^64,77,82^ were synthesized by Twist Bioscience. The DNA fragments were flanked with biobrick prefix and suffix for PCR and Gibson assembly purposes. The fragment also contained two SapI restriction sites for insertion of the gene of interest after the intein, and two BsaI restriction sites to transfer the fusion gene cassette to a new backbone with a selected promoter (see *Supplementary Figure S1*). The synthesized DNA fragments were amplified by PCR with primers MFL 25 & 26 and subsequently purified (see primer list in *Supplemental Table S1*). A pUC8 backbone was amplified by PCR with primers MFL 334 & 335, containing complementary overhangs to the synthesized DNA fragments. Each mini-intein DNA fragment was assembled onto the pUC8 backbone by Gibson assembly^83^ (see *Supplementary Figure S1a*). Then, 10 μL of each assembly mixture was chemically transformed into competent *E. coli* (heat shock 42°C, 45s) and cells were plated on LB-agar plates containing 100 μg/mL ampicillin. Correct insertion was assessed by colony PCR with primers MFL 25 & 26. Positive clones were grown overnight at 37°C in 5 mL LB medium (10 g/l tryptone, 5 g/l yeast extract, and 5 g/l NaCl) containing 100 μg/mL ampicillin, the respective plasmids were purified by Miniprep and the sequence was confirmed by Sanger DNA sequencing.

Three genes of interest were used to test the intein standard expression system, *sfGFP*, *mCherry* and the chloramphenicol acetyltransferase gene *cat*. *sfGFP* was PCR amplified in two parts from the biobrick (BBa_I746916) with primers MFL 355 & 356 and MFL 357 & 132 respectively, in order to eliminate an internal BbsI restriction site and insert a C-terminal 6-His tag onto *sfGFP*. The *E. coli* codon optimized *mCherry* gene was synthesized by Twist Bioscience and all internal BbsI restriction sites were removed by using synonymous codons. The *mCherry* gene was amplified with primers MFL 351 & 354. The *cat* gene was amplified from the pXMJ19 plasmid with primers MFL 359 & 318. The PCR amplifications conferred each gene with upstream and downstream SapI restriction sites, with CCA and TAA scars respectively, complementary to the SapI scars of the intein DNA fragments. Each gene was assembled onto the pUC8-DnaB and pUC8-DnaX plasmids by cycles of SapI restriction (2 min, 37°C) and ligation with T4 ligase (2 min, 16°C) as presented in the Start-Stop assembly method^84^ (see *Supplementary Figure S1b*). 10 μL of each Start-Stop assembly mixture was chemically transformed into *E. coli* and cells were plated on LB-agar plates containing 100 μg/mL ampicillin. Colony PCR with primers MFL 25 & 26 was performed to check the correct insertion of the genes of interest. Positive clones were grown overnight at 37°C in 5 mL LB containing 100 μg/mL ampicillin and the respective plasmids were purified and sequenced. The resulting plasmids pUC8-DnaB-GOI and pUC8-DnaX-GOI were used as template for the amplification of the standard cassettes.

The *GumM* gene and the *CBM73* gene were PCR amplified from *Kozakia baliensis SR745* genomic DNA^74^ and from plasmid pNIC-CH-CBM73^73^ using primers MFL 1042 to 1044 (see *Supplemental Table S1*). The backbone was amplified from Lv1-P7-DnaB-GFP with primers 1040 and 1041 to excise GFP and be used as template in start/stop assembly cloning. The *GumM* and *CBM73* gene were cloned onto the backbone as presented above using the Start-Stop assembly method^84^, transformed into E. coli Dh5α, verified by colony PCR. Positive clones were grown overnight at 37°C in LB medium and their plasmid was purified by miniprep and sequence verified by Sanger sequencing. Once verified, the respective plasmids were transferred to E. coli BL21 strains by heat shock and positive clones were grown in 5mL overnight as precultures for protein production.

### Creation of promoter library and promoter selection

To create a promoter library adapted to *DnaB* and *DnaX* gene sequence, we use the GeneEE method^6285^ that utilizes 200-nucleotide long DNA fragment of random composition to generate gene-tailored promoters and 5’UTRs. In brief, a single stranded DNA fragment of 200 random nucleotides (200N), synthesized by Integrated DNA Technology (Louvain, Belgium), was amplified by PCR with primer MFL 25 & 26. The resulting double stranded DNA fragment contained BsaI restriction sites with an upstream 5’-TGCC-3’ scar and a downstream 5’-NATG-3’ scar. The pUC8-DnaB-GFP and pUC8-DnaX-GFP were used as templates for PCR amplification of the *DnaB-GFP* and *DnaX-GFP* gene fragments (primers MFL 25 & 26) followed by treatment with DpnI overnight at 37°C. The *DnaB-GFP* and *DnaX-GFP* gene fragments both contained an upstream 5’-AATG-3’ BsaI scar, but a different downstream BsaI scar, 5’-AGTT-3’ for *DnaB-GFP* and 5’-TCAA-3’ for *DnaX-GFP*. The random 200N DNA fragment and the intein-GFP fragment were assembled on pUC19 backbones with corresponding BsaI scars by a 3-piece Golden Gate assembly^86^ (see *Supplementary Figure S1c*). Two *E. coli* transformations were performed for each Golden Gate mixture and cells were plated on LB-agar plates containing 100 μg/mL ampicillin. For both *DnaB-GFP* and *DnaX-GFP*, colonies seemingly displaying green fluorescence under UV light were picked and grown overnight at 37°C at 800 rpm into two 96-well plates containing LB supplemented with 100 μg/mL ampicillin.

Cells were transferred into black 96-well plates with transparent bottom (Thermo Scientific) and fluorescence was measured with an Infinite M200 Pro TECAN fluorimeter (Noax Lab AS). GFP was excited with a wavelength of 488 nm and fluorescence emission was detected at 526 nm. The fluorescence values were normalized by the OD_600_ of the corresponding well. For both *DnaB-GFP* and *DnaX-GFP*, 10 strains representing different expression levels were selected and grown in triplicates overnight at 37°C with 800 rpm agitation in 96-well plates with LB supplemented with 100 μg/mL ampicillin. Strains displaying inconsistent fluorescence values when replicated were abandoned. Seven strains for *DnaB-GFP* and eight for *DnaX-GFP* were conserved to represent the expression range and each GES was sequenced (see *Supplementary Table S3*).

### Promoter transfer to other gene cassettes

In order to assess whether the strength of GES remains equivalent when other genes are fused to the inteins, we tested the *mCherry* gene and the chloramphenicol resistance (CmR) gene *cat*. Each GES selected for *DnaB-GFP* and *DnaX-GFP* was extracted by PCR with primers MFL 330 & 331 and MFL 332 & 333, respectively. These primers contain BsaI recognition sites that confer an upstream 5’-TGCC-3’ scar and a downstream 5’-AATG-3’ scar on each side of the GES sequences. The *DnaB/X-mCherry* and *DnaB/X-cat* cassettes were amplified by PCR with primer MFL 25 & 26 from the pUC8-Intein-GOI plasmids described above, and digested with DpnI overnight at 37°C. As for *GFP*, the extracted GES sequences and the Intein-GOI cassettes were assembled on pUC19 backbones with corresponding BsaI scars by a 3-piece Golden Gate assembly^86^ (see *Supplementary Figure S1c*). Each Golden Gate mixture was transformed into *E. coli* and plated on LB-agar plates containing 100 μg/mL ampicillin for *mCherry* constructs and 10 μg/mL chloramphenicol for *cat* constructs. Colonies selected on chloramphenicol and visibly red colonies under UV light were grown in triplicates overnight at 37°C with 800 rpm agitation in 96-well plates in LB media supplemented with the appropriate antibiotic.

Cells carrying *DnaB/X-mCherry* plasmids were transferred into black 96-well plates with transparent bottom to measure their respective fluorescence with an Infinite M200 Pro TECAN fluorimeter. The fluorescence of mCherry was monitored with a wavelength couple of 576/610nm and fluorescence values were normalized by the OD_600_ of the corresponding well. Cells displaying fluorescence levels deviating from expectations based on GFP fluorescence levels were grown overnight in 5 mL LB supplemented with 100 μg/mL ampicillin for plasmid purification and sequencing.

Cells carrying *DnaB/X-cat* plasmids grown in a 96-well plate were diluted 100 times into a new 96-well plate containing LB supplemented with 10 μg/mL chloramphenicol. The dilutions were stamped on 15 cm diameter LB-agar plates containing increasing chloramphenicol concentrations (10, 30, 50, 75, 100, 150, 200, 250, 300, 350, 400 and 500 μg/mL) using a 96-pin replicator. Cells were incubated overnight at 37°C and photographed with a Canon camera EOS M.

### Coupling of intein cassettes

The gene cassettes described above with *DnaB-GOI* and *DnaX-GOI* were assembled on pUC19 backbones from the previously described pathway assembly method (Fages-Lartaud *et al*^67^). The principles of the method that are utilized in this study are described hereafter. In the first level, promoter libraries or selected promoters are assembled with a single gene on a respective pUC19 backbone (Lv1 plasmids) (see *Supplementary Figure S1c*). Each Lv1 plasmid contains an upstream and a downstream BbsI site. Here, the Lv1 carrying *DnaB-GOIs* contains and upstream 5’-TGCC-3’ BbsI scar and a downstream 5’-AGTT-3’ BbsI scar; and the Lv1 carrying *DnaX-GOIs* contains and upstream 5’-AGTT-3’ BbsI scar and a downstream 5’-TCAA-3’ BbsI scar. The *DnaB-GOI* and *DnaX-GOI* cassettes can be coupled on a Lv2 plasmid containing matching 5’-TGCC-3’ and 5’-TCAA-3’ BbsI scars that is selectable on chloramphenicol (see *Supplementary Figure S1d*).

Different combinations of *DnaB-mCherry* and *DnaX-GFP* with various GES intensities were tested. The respective Lv1 plasmids were mixed together with a Lv2 backbone and subject to 50 Golden Gate-like assembly cycles for BbsI restriction (2 min, 37°C) and T4 ligase ligation (2 min, 16°C)^87 67^. The resulting assembly mixtures were transformed into *E. coli* cells that were plated on LB-agar plates containing 30 μg/mL chloramphenicol and incubated overnight at 37°C. Positive clones were grown in triplicates, overnight at 37°C with 800 rpm agitation, in 96-well plates in LB supplemented with 30 μg/mL chloramphenicol. Cells were transferred into black 96-well plates with transparent bottom to measure the fluorescence of GFP and mCherry as described previously. GES sequences were sequenced in constructs that resulted in discrepant fluorescence intensities compared to the expectations based on fluorescence measurements of the single-gene constructs.

### Fluorescence microscopy

The strains carrying the weakest single-gene GES (P1) and the weakest paired-gene GES (P2) were analyzed by fluorescence microscopy to confirm protein expression and cellular co-localization. One microliter of overnight culture was analyzed with an inverted microscope (Zeiss Axio Observer.Z1, 14 2.3.64.0) possessing a 20x air objective (NA 0.8). The GFP and mCherry filters were applied to measure the fluorescence of both proteins. Image processing was performed with Zeiss image analysis software (2.3.64.0).

### Intein inactivation for the assessment of protein cleavage

In order to verify the correct C-terminal cleavage of intein, we compared intein cleavage with inactive, C-terminal truncated variants of *DnaB* and *DnaX*. The plasmids carrying a strong GES expressing *DnaB-GFP* and *DnaX-GFP* were used for the analysis of C-terminal cleavage. The Lv1-P7-*DnaB-GFP* and Lv1-P8-*DnaX-GFP* were amplified by PCR with primers MFL 633 & 132 and MFL 634 & 132 respectively. The PCR creates a C-terminal truncation of the *DnaB* and *DnaX* genes and primers contain SapI sites necessary for Start/Stop assembly. PCR products were digested with DpnI overnight at 37°C and subsequently purified. The single fragments were subject to 50 cycles of SapI digestion (2 min, 37°C) and ligation with T4 ligase (2 min, 16°C) as described before. A volume of 10 μL of each Start/Stop assembly mixture was transformed into *E. coli* cells. Transformants were selected on LB-agar plates containing 100 μg/mL ampicillin and incubated overnight at 37°C. Several clones were grown overnight at 37°C in 5 mL of LB medium containing 100 μg/mL ampicillin and their respective plasmids were purified and sequenced. The *sfGFP* gene contains a C-terminal 6-His tag suitable for protein purification.

### Protein production and purification

The Lv1-P7-*DnaB-GFP* and Lv1-P8-*DnaX-GFP* plasmids and their mutated versions were transformed into *E. coli* BL21 for protein production. Positive clones were grown overnight at 37°C in 5 mL LB medium supplemented with 100 μg/mL ampicillin. A volume of 2 mL of overnight culture was used to inoculate conical Erlenmeyer flasks containing 200 mL of the same medium. The cultures were incubated in a shaking incubator at 30°C and 225 rpm for 24 hours.

For larger scale protein production, the Lv1-P7-DnaB -GFP, -CBM73 and -GumM plasmids were transformed into *E. coli* BL21 for protein production. Positive clones were grown overnight at 30°C and 225 rpm in 5 mL LB medium (10 g/l tryptone, 5 g/l yeast extract, and 5 g/l NaCl) supplemented with 100 μg/mL ampicillin. These cultures were used to inoculate 500 mL 2xLB medium (20 g/l tryptone, 10 g/l yeast extract, and 5 g/l NaCl) supplemented with 100 μg/mL ampicillin. The cultures were incubated at 22°C or 37°C in a LEX-24 Bioreactor (Harbinger Biotechnology) for 24 hours.

Cells were harvested by centrifugation (4500 × g, 10 min) followed by cell lysis using pulsed sonication in 10 mL lysis buffer (50 mM Tris-HCl, 50 mM NaCl, 0.05 % Triton X-100, pH 8.0) supplemented with 1 tablet EDTA-free cOmplete ULTRA protease inhibitor (Roche). Cell debris were removed by centrifugation (30,000 × g, 30 min). The supernatant was sterilized by filtration through a 0.2 μm Nalgene sterile vacuum filter unit (ThermoFischer). Lysates were used for Western blot analysis after SDS-PAGE migration (see below). Samples for LC-MS analysis were purified on Nickel-sepharose column and analyzed on SDS-PAGE (see below).

### Protein purification

Lysates were purified using a 1 mL HisTrap HP Ni-sepharose column (GE Healthcare Life Sciences) connected to an ÄKTA pure protein purification system (Cytiva). The column was equilibrated with 5 CV Buffer A (50 mM Tris-HCl, 300 mM NaCl, pH 8.0) before loading the supernatant onto the column. Impurities were removed by washing with Buffer A for 10 CV. His-tagged proteins were eluted using a gradient of 0-100% Buffer B (50 mM Tris-HCl, 300 mM NaCl, 400 mM imidazole, pH 8.0) over 40 CV. All steps were performed using a flow rate of 1 mL/min. GFP-containing fractions containing were identified using absorbance at 488 nm.

### Protein analysis

SDS-PAGE gels were run under denaturing conditions using SurePAGE™ Bis-Tris 12% gels (GenScript) and Tris-MES-SDS Running buffer (GenScript). The gel used for Western blotting was not stained, whereas all other gels were stained using the eStain™ L1 protein staining system (GenScript). Precision Plus Protein™ standards (Bio-Rad Laboratories) or PAGE-MASTER Protein Standard Plus (GenScript), 5 μL in both cases, were used for the identification of target proteins.The gels were analyzed using the ImageJ software. Protein yields were estimated from the amounts of protein in the gels, *M_prot_*, by using the following formula,

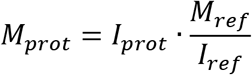

where *I_ref_* corresponds to the sum of intensities of all nine bands in the PAGE-MASTER Protein Standard Plus standard lane, *M_ref_* is the total amount of protein applied in the standard lane (equal to 8 μg), and *I_prot_* is the intensity of the band of the protein of interest.

For Western blot, proteins were transferred from the unstained gel to polyvinylidene fluoride membranes using the Trans-Blot turbo transfer system (Bio-Rad Laboratories). An iBind™ Flex system (Invitrogen/ThermoFischer) was used for blocking and antibody incubation according to the manufacturer’s protocol. The primary antibody (6xHis tag monoclonal antibody; Invitrogen/ThermoFischer), secondary antibody (Polyclonal rabbit anti-mouse immunoglobulins/horseradish peroxidase; Dako/Agilent) and iBind™ Flex solutions were added to their corresponding reservoirs in the iBind™ Flex system. Following incubation for 1 h at room temperature, membranes were rinsed in water prior to immunodetection with 3,3’,5,5’-tetramethylbenzidine (Sigma-Aldrich/Merck).

### Sample Preparation for LC-MS

Please note that (unless otherwise specified): volumes are to cover the gel bands, also liquids are removed after each incubation. The excised gel bands of interest were cut in smaller pieces (3-5 mm3) and were de-stained by incubation for 15 minutes in 50 mM ammonium bicarbonate (ABC), 50% methanol. Then, they were shrunk with acetonitrile for 15 minutes. The samples were reduced by 10 mM DTT in 25 mM ABC at 56°C for 45 minutes, alkylated by 55 mM iodoacetamide in 25 mM ABC at room temperature in the dark for 45 minutes, washed with 50 mM ABC, 50% methanol for 10 minutes and shrunk with acetonitrile. Then, 12.5 ng/μL trypsin in 50 mM ABC was added to the gel pieces and incubated for 30 minutes on ice. After removal of the liquid, 50 mM ABC was added, and samples were digested by trypsin at 37°C overnight. Peptides were collected and dried in a vacuum concentrator at room temperature. Dried peptides were reconstituted in 50 μL 0.1% formic acid in water and shaken at 6 degrees Celsius at 900 rpm for 1.5 hour. Samples were centrifuged at 16,000 × g for 10 minutes and 40 μL of the supernatants were transferred to MS-vials for LC-MS analysis.

### LC-MS analysis

LC-MS analysis was performed on an EASY-nLC 1200 UPLC system (Thermo Scientific) interfaced with an Q Exactive mass spectrometer (Thermo Scientific) via a Nanospray Flex ion source (Thermo Scientific). Peptides were injected onto an Acclaim PepMap100 C18 trap column (75 μm i.d., 2 cm long, 3 μm, 100 Å, Thermo Scientific) and further separated on an Acclaim PepMap100 C18 analytical column (75 μm i.d., 50 cm long, 2 μm, 100 Å, Thermo Scientific) using a 120-minute multi-step gradient (90 min 5%-40% B, 15 min 40%-100% B, 15 min at 100% B; where B is 0.1 % formic acid and 80% CH_3_CN and A is 0.1 % formic acid) at 250 nl/min flow. Peptides were analyzed in positive ion mode under data-dependent acquisition using the following parameters: Electrospray voltage 1.9 kV, HCD fragmentation with normalized collision energy 25. Each MS scan (200 to 2000 m/z, 2 m/z isolation width, profile) was acquired at a resolution of 70,000 FWHM in the Orbitrap analyzer, followed by MS/MS scans at resolution 17,500 (2 m/z isolation width, centroid) triggered for the 8 most intense ions, with a 60 s dynamic exclusion and analyzed in the Orbitrap analyzer. Charge exclusion was set to unassigned, 1, >5.

### Processing of LC-MS Data

Proteins were identified by processing the LC-MS data using Thermo Scientific Proteome Discoverer (Thermo Scientific) version 2.5. The following search parameters were used: enzyme specified as trypsin with maximum two missed cleavages allowed; acetylation of protein N-terminus with methionine loss, oxidation of methionine, and deamidation of asparagine/glutamine were considered as dynamic and carbamidomethylation of cysteine as static post-translational modifications; precursor mass tolerance of 10 parts per million with a fragment mass tolerance of 0.02 Da. Sequest HT node queried the raw files against sequences for expected proteins (DnaB GFP, DnaX GFP, BM GFP, GFP C-terminal) and a common LC-MS contaminants database. For downstream analysis of these peptide-spectrum matches (PSMs), for protein and peptide identifications the PSM FDR was set to 1% and as high and 5% as medium confidence, thus only unique peptides with these confidence thresholds were used for final protein group identification and to label the level of confidence, respectively.

## Notes

### Competing Interest Statement

The authors have declared no competing interest.

### Summary of Updates

The manuscript has been complemented with data regarding intein cleavage condition and the production of difficult-to-express protein using SIGER systems

