## Supplementary data for "Standard Intein Gene Expression Ramps (SIGER) for protein-independent expression control"

### Supplementary Figures and Tables

### Standard Intein Gene Expression Ramps (SIGER) for controlling enzymatic ratios *in vivo*

Fages-Lartaud *et al. 2023*

### Table of Content

Supplementary Table S1: Primer list……………………………………………………………………………p3

Supplementary Table S2: Intein sequences…………………………………………………………………..p4

Supplementary Table S3: GES sequences for DnaB and DnaX……………………………………….p5

Supplementary Figure S1: Scheme of the cloning process……………………………………………..p6

Supplementary Figure S2: Chloramphenicol gradient for *DnaB-cat* and *DnaX-cat*………….p7

Supplementary Figure S6. Purification of GFP from cultures grown at 22 °C and 37 °C …….p11

#### Supplementary Table S1: Primer list

| **Name** | Comment | Sequence |
|---|---|---|
| **MFL 025** | Biobrick prefix F | GAATTCGCGGCCGCTTCTAGAG |
| **MFL 026** | Biobrick suffix R | CTGCAGCGGCCGCTACTAGTA |
| **MFL 334** | pUC8- Biobrick-F | TACTAGTAGCGGCCGCTGCAGTTGGCACTGGCCGTCGTTTTACAAC |
| **MFL 335** | pUC8- Biobrick-R | TCTAGAAGCGGCCGCGAATTCGCTTCCTCGCTCACTGACTCGC |
| **MFL 318** | cat-SapI-R | GAACTCGCTCTTCATTATTCAAATGTAGCACCTGAAGTCAGCC |
| **MFL 351** | mCh-SapI-F | GAACTCGCTCTTCACCAGTGAGCAAGGGCGAGGAGGAT |
| **MFL 354** | mCh-SapI-R | GAACTCGCTCTTCATTACTTGTACAGCTCGTCCATGCC |
| **MFL 355** | GFP-SapI-F | GAACTCGCTCTTCACCACGTAAAGGCGAGGAGCTG |
| **MFL 356** | GFP-SapI-P1-R | GAACTCGCTCTTCCATCTTCTTTAAAGTCAATGCCT |
| **MFL 357** | GFP-SapI-P2-F | GAACTCGCTCTTCAGATGGCAATATCCTGGGCCATAA |
| **MFL 132** | sfGFP -SapI-R | AGGATTGCTCTTCATTAGTGGTGGTGGTGGTGGTGTTTATACAGTTCGTCCATACCGTG |
| **MFL 359** | cat-SapI-F | GAACTCGCTCTTCACCAGAGAAAAAAATCACTGGATATACCACCG |
| **MFL 360** | DnaB/X-SapI-R | AGGATTGCTCTTCATGGTCCCCTAGAGTT |
| **MFL 263** | Lv1-BsaI-TGCC-R | AGAGTTGGTCTCAGGCAGGGTCTTCGATCGTGCATCAG |
| **MFL 264** | Lv1-BsaI-AGTT-R | AGAGTTGGTCTCAAACTGGGTCTTCGATCGTGCATCAG |
| **MFL 266** | Lv1-BsaI-AGTT-F | AGAGTAGGTCTCAAGTTCCGTCTTCAAGCGAGAACGC |
| **MFL 268** | Lv1-BsaI-TCAA-F | AGAGTAGGTCTCATCAACCGTCTTCAAGCGAGAACGC |
| **MFL 330** | DnaB-200N-extract-R | ACTATCGCCAGAGATAGCTCCGGGGTCTCGCATT |
| **MFL 331** | Lv1-DnaB-200N-F | GCCCTTTCCTGATGCACGATCGGTCTCCTGCC |
| **MFL 332** | DnaX-200N-extract-R | CCTTTTATTTGGGGATTGTCAATGGACATTAGGTCTCTCATT |
| **MFL 333** | Lv1-DnaX-200N-F | CCTTTCCTGATGCACGATCGAAGAGGTCTCTTGCC |
| **MFL 633** | DnaB-mut-R | AGTGTGGCTCTTCGTGGTCCTGGCACAGTCAAATCAAA |
| **MFL 634** | DnaX-mut-R | AGTGTGGCTCTTCGTGGGATTATCTTCCACCTCCAGGTC |
| **MFL 1040** | DnaB-SapI-R | AGGATTGCTCTTCATGGTCCCCTAGAGTTGTGTACAAT |
| **MFL 1040** | DnaB-SapI-F | GAACTCGCTCTTCATAAATCTGCGATTCTGATAACAAACTAGC |
| **MFL 1041** | X75-SapI-F | GAACTCGCTCTTCACCAGGCAATTGTATCAGTCCGGTGT |
| **MFL 1042** | X75-Histag-SapI-R | GAACTCGCTCTTCATTAATGGTGGTGATGATGGTG |
| **MFL 1043** | GumM-SapI-F | GAACTCGCTCTTCACCATTTAATGTCATGGGGGTGCGCC |
| **MFL 1044** | GumM-His-SapI-R | GAACTCGCTCTTCATTAATGGTGGTGATGATGGTGGCCGGCAGCGTGGTCGACCAAG |

#### Supplementary Table S2: Intein sequences

Nucleotide and amino acid sequences of *DnaB* and *DnaX*. The N-terminal asparagine responsible for cyclization and autocatalytic release of the POI is highlighted in blue. The linker peptide that connects the inteins and the POIs in highlighted in gray. The SapI scar CCA that matches the GOI is underlined in purple.

| | |
|---|---|
| **DnaB** | atgcgcgagtccggagctatctctggcgatagtctgatcagcctggctagcacaggaaaaagagtttctattaaagatttgttagatga<br>aaaagattttgaaatatgggcaattaatgaacagacgatgaagctagaatcagctaaagttagtcgtgtattttgtactggcaaaaag<br>ctagtttatattctaaaaactcgactaggtagaactatcaaggcaacagcaaatcatagatttttaactattgatggttggaaaagatt<br>agatgagctatctttaaaagagcatattgctctaccccgtaaactagaaagctcctctttacaattgtcaccagaaatagaaaagttgt<br>ctcagagtgatatttactgggactccatcgtttctattacggagactggagtcgaagaggtttttgatttgactgtgccaggaccacata<br>actttgtcgcgaatgacatcattgtacacaactctaggggacca |
| | MRESGAISGDSLISLASTGKRVSIKDLLDEKDFEIWAINEQTMKLESAKVSRVFCTGKKLVYILKTRLGRTIKAT<br>ANHRFLTIDGWKRLDELSLKEHIALPRKLESSSLQLSPEIEKLSQSDIYWDSIVSITETGVEEVFDLTVPGPHNF<br>VANDIIVHNSRGP |
| **DnaX** | atgaatggcttaatgtccattgacaatccccaaataaaagggcgagaagttctgagctacaacgaaactctacagcaatgggaatat<br>aaaaaagttttaagatggcttgacagaggcgaaaagcaaacattgtctattaagacaaaaaattctacagtacggtgtacggctaac<br>catttaatcagaactgaacaaggatggacgagagcggaaaacatcactcccggaatgaagatactatcccctgcctctgcctctggac<br>atcaccaccatcaccatggaggctccggctcctccccgcaatggcatacaaatttcgaggaagttgagtccgtcactaagggtcaagt<br>ggaaaaagtttatgacctggaggtggaagataatcacaattttgttgccaatggcttattagtccataactctaggggacca |
| | MNGLMSIDNPQIKGREVLSYNETLQQWEYKKVLRWLDRGEKQTLSIKTKNSTVRCTANHLIRTEQGWTRA<br>ENITPGMKILSPASASGHHHHHHGGSGSSPQWHTNFEEVESVTKGQVEKVYDLEVEDNHNFVANGLLVH<br>NSRGP |

#### Supplementary Table S3: GES sequences for DnaB and DnaX

Promoter motifs similar to the -35 and -10 boxes, TTGACA and TATAAT respectively, are highlighted in blue. SD-like sequences similar to GGAGGA and close to the start codon are highlighted in orange.

| | Number | GES Sequence |
|---|---|---|
| DnaB | B_P1 | GTTAGGGAATTTTTTAACATGTTAGGATGTCTAAGTATTTCGACCTTTTTTGAACCTACCTTGGGAGATG<br>TGGTATGCGTACCACATGATACTTTTAAGTCGTCGCCACATAGCGGTTCTCTGTCGATTTGCGGACGTGA<br>ACTTCGTTCGTTATAGTAGGCAGTATGATTTTATCGGCTGCAGAATGTA |
| | B_P2a | AAAGTCTTCTTGATCCGCACGCCGTTAATCAGTATCAGGTTGTCTATGGGATGTAAAAGTGTTATATATT<br>GGTCAGTTCGGGCTCAAGTTGATTTATATATTCCACTCTTATAGAGTGGTAATCGCGGACGGGCGCTCG<br>TGCCATAATAATCACGATGGCGGGTCAGGGCCAATGGCTAGCGCGAGGAGGATGCGGAGTT |
| | B_P2b | ACACTACAAGGTTTCTTTATAAATTTCCTACTAGCAATTTCATAGTTGTTCGATTGGTTGTCGTTACTAGA<br>GGCATTATCATGTCTCAACTTATGGCGATTTAACGTCATTGGCCTCGAGGCTTGACGTAGATTCGCCTCC<br>TAGTTGTACCGGAGGGGACTTAGGGTGGTTTGGTGGGAGAGGTGTATTCCCTTTATTTA |
| | B_P3 | AGTGCACCGTCTGGTATCTAAATATGTTTATCTTTGGAGTGTTGGCAGGAACGTTGTGTCCAGCCTTAAA<br>ATGGTTTATTATGGTGGTATACTTATATTCACATGTGAAAAGGGCAATGCGATGTTAAGATGCGACTTG<br>CAACAGCAGGATGTTCCTGTAGTCCCCCGGTCCAGCAAGGAATGGTGGTCCATTCCCACTA |
| | B_P4 | GTCTCGTGATGGCCGCTGTGTTAAGATCTACAGAGAGATTCAGTCGGACTGAAGTACAGTCAGAAGTTT<br>GGAAGTGGACAGTTAGTAAGATGTTGCACTTATTCCGGAGGCCAGGGTACAATATGGCAAGCGCCCAC<br>TCCCTTGCCCGAACTGGCAGTTCGTTTTGATGCTCTCAGGTCGGACAAGGTGTGGGATGCCTA |
| | B_P5 | CGTTTGCATTTACGTGTAAGTCTGTGCTAACCAAGGCTCGTTTCCATCAGTGGCCGAGTAGATTATTATC<br>TCGAATACTTGTCAGCTATTAGAGGTTGAGGTAAAGTGTCGTGGTGTCGGAGCGAAGGGATTTTAGGG<br>GCTCTGGGCTGGTGGAATATGCAGTACGGCAAGCGAGACGTTTGGGCGTTACCATCGTTAGTA |
| | B_P6 | AATTCTACTAGATGGACTTACGTAATAGCCGTTTTTGCAGCGTCGCTAGGAAATAGATTTCCTCGTGTGC<br>TTTATCGCACGTAATGGTGCTCGGTGTTCCGGTCACCGTGCAGATTTCCGCCCTGGAGATAGGATTTAA<br>CTTGCTTCTCTCCTCCGTTTTTTGTAAGTTAGTTGTGATTTCTGAAGGCCTTGTGTCTTCA |
| | B_P7 | ACGCAACCCAAGTAAAACTCGATTTAGCCTCAGGCGCGTGCTATAGTTACGTAGTGTTCGGAGTATGCA<br>GGCCTGTTTCAGACATCGAGAGTAAAGTGACAGGGTTGAGGGTAACCTGTCGACTGGAAGGGCAACC<br>ATGGCTGTCGTTCCGATTGCGGATGGATAGAAGTGGCGGCA |
| DnaX | X_P1 | GTGTGATGCCATGGCGCGAAGAATATTGAAAGTGTAGTTTTGACATACTATTGTCTTATGTGATTTCATA<br>CTTTCGATCTAGTTAGCGGACTCTCCTATCTTTGGTTTCAAATGTAGTTTATGAGCGCTACTTGTAATTCA<br>CGTCATCTGTTCCACCGCTTGTAAGCAGGGATGGCGAGCTGTGATTCACCAGTGGA |
| | X_P2 | AACAAGATAGAGATATAAATTGCATTGTCACTTTATGTGTATGATATTTGCTATATGCATATAGAATTCG<br>AAGGTAGGTGGTGCTTGCCGAATGAGAAAGTCTTTTATATTATGCGTTCGTATAGGAGATACGATAGCT<br>GGTGCCGGATATTAATTGTGCAGCACGGTTCTAAGACTACCGGAGCGTGCACCCACGGCTT |
| | X_P3 | ATTTATTGCGGAATTTAGTATACTAGCCCTCTATAGGCTTATGTTGCATGTGGGTTAATCGTCGCTTACC<br>ATCTAAGTGGATAGTTACACTCTCCCAGTCACACAATTCTGACTAGAAAGACGTATTAATCGCTTCGGAT<br>TGGTTTACTTTGAGTTAGTACCACTTTGGGTTGGTTGTTCTGGTATTTGGGTTCTTGGGAG |
| | X_P4a | ATGTGTTCGTACCCGTCTTAAAATTTACATATTATGAGTAAGGGAACAGGTTACTAGACCGATACCGCC<br>ATAAGTTGAGTTCTCTTACATCGCGTGGACATGTTGGAATAGGGACAATTTGTCGACACCTGACATCTTA<br>AAAGCTAGAATCCTATTGTAAGCGAAACAGCTGAATTTACTCTAACTGTAGGACTTATTAA |
| | X_P4b | AACAAGATAGAGATATAAATTGCATTGTCACTTTATGTGTATGATATTTGCTATATGCATATAGAATTCG<br>AAGGTAGGTGGTGCTTGCCGAATGAGAAAGTCTTTTATATTATGCGTTCGTATAGGAGATACGATAGCT<br>GGTGCCGGATATTAATTGTGCAGCACGGTTCTAAGACTACCGGAGCGTGCACCCACGGCTT |
| | X_P5 | TTGCATTCTATATAGCGTCTAGTTAAACTAATTGGACACTCGCCGAGCGTCTTCATGTACTACGTAGATG<br>TATTGACACTCGCGCGAGTATGAACATATGATGTTCGATATAGGGTGCGTCTGAACACCTACCTTTTTAG<br>GTTCTTACGTTAAAGATACCTTGGTGGCTGCTACCGCGATCGGGCAGGTAATCCTTGAAA |
| | X_P6 | CACGTCTCGGGATGTAGTGATGTCGGTCTTACTGCATGGTGGTTATCAACCTGGTGTTGTGGGGGGTAT<br>TGGCTCTAGTCTGTACTTCTTGGTCATATGTAGCTCTAGCCGCGCTAGTTGTACGTTGACAATGGTTCGT<br>ATGATTAGTAATATAGCATCTTGATGATGGCATAGAGGAACTGCATAGACACGGCGTGGAA |
| | X_P7 | TTCAAGGATGTCGTAGTGACTGGTGATGCGTTCGTTTATGAGCTATTGAGGTAACGATCAGGGTTTATT<br>TTTATAAGGTTCTAGTAAGCGTATTGCAGAAATGGAACCTTGTGTTGTTTGTGTACATTGACTAGGTTCC<br>CAATGTGCGATATGGTTAGGAGCATAAGAATCACGCGTAGCTGCTTAGAGATGAGTGTAT |
| | X_P8 | CCCAAAGGAAGAGAGAACTTAAGTAAATGGCCATCATAGGATCGCTGGGATTGTGTATCGATCCAGAG<br>TCTAGTAGTAATTTGATTTTTTCGAGTTTAAAGACTATAAGAAGACGCTTGACCCGTGTGTATGATCATA<br>GTGTAGTGCGGGTAGCAAAGTTACGATATGGTGGGGAGGATTTCGATATGAAGGCTAGTACA |

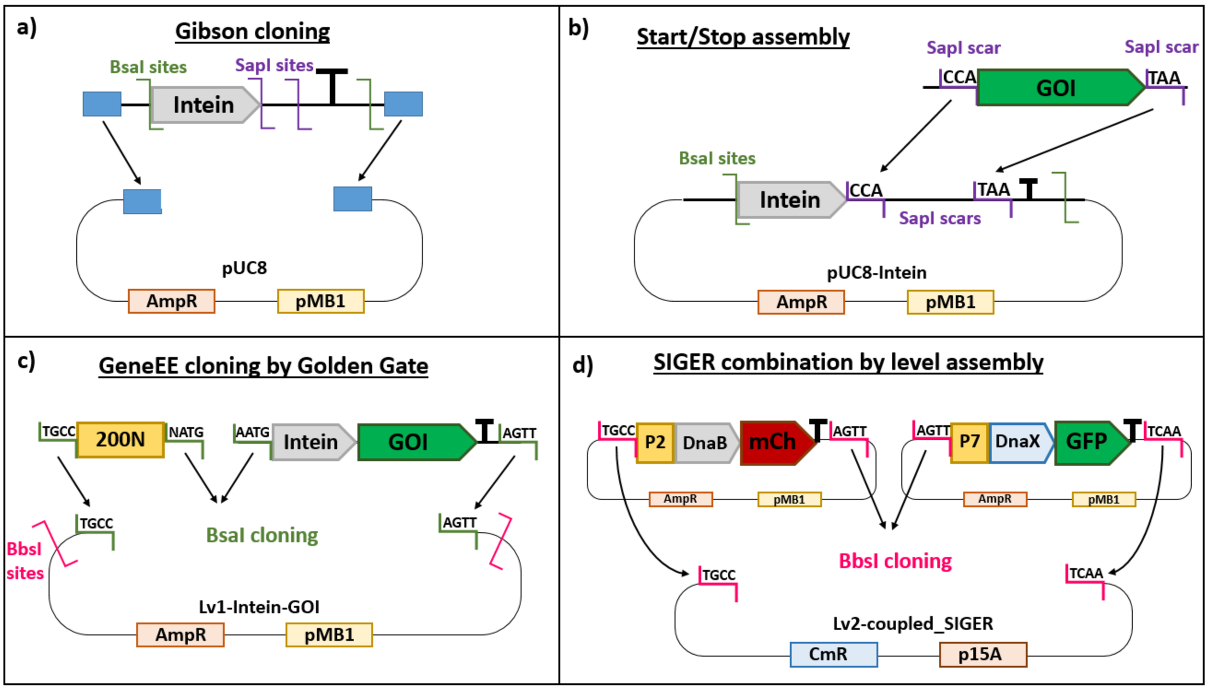

#### Supplementary Figure S1: Scheme of the cloning process

a) Gibson cloning of the synthesized Intein cassettes onto a pUC8 vector containing the complementary Biobrick sequences. The Biobrick sequences are not used for restriction enzymes but only for sequence complementarity for the Gibson assembly. The Intein cassette contains a BsaI site with NATG scar at the intein gene-start, and another BsaI site with AGTT or TCAA scars for *DnaB* and *DnaX* respectively after the terminators (BBa_1006_U10 for *DnaB* and rrnBT1 + T7Te for *DnaX*). The intein genes are followed by SapI sites with CCA and TAA scars for GOI insertion.

b) Insertion of the GOIs by Start/Stop-like assembly. SapI sites were included on the PCR forward and reverse primers of *sfGFP*, *mCherry* and *cat* to provide CCA and TAA SapI scars on each side of the genes. Start/Stop assembly cycle of SapI restriction (37°C) and ligation with T4 ligase (16°C) inserted seamlessly the GOIs after each intein.

c) The GES library was created by inserting the 200N random DNA fragment upstream the *DnaB-GFP* and *DnaX-GFP* cassettes onto a Lv1 plasmid using a 3-piece Golden Gate assembly. The three BsaI scars match to assemble seamlessly 200N to the inteins onto the plasmid. The *DnaB-GFP* construct was assembled on a Lv1 with an upstream TGCC scar and a downstream AGTT scar; and *DnaX-GFP* was assembled on a Lv1 with an upstream AGTT scar and a downstream TCAA scar. The Lv1 plasmids contain BbsI sites that leave the respective upstream and downstream scars.

d) The pathway level assembly was used to assemble SIGER systems on a Lv2 plasmid. The Lv1 plasmids carrying the *DnaB-mCherry* and *DnaX-GFP* cassettes with the selected GES were mixed with a Lv2 plasmid containing a chloramphenicol resistance gene. The BbsI restriction/T4 ligation cloning cycles assemble the two SIGER systems onto the Lv2 plasmid due to the complementarity between the BbsI scars.

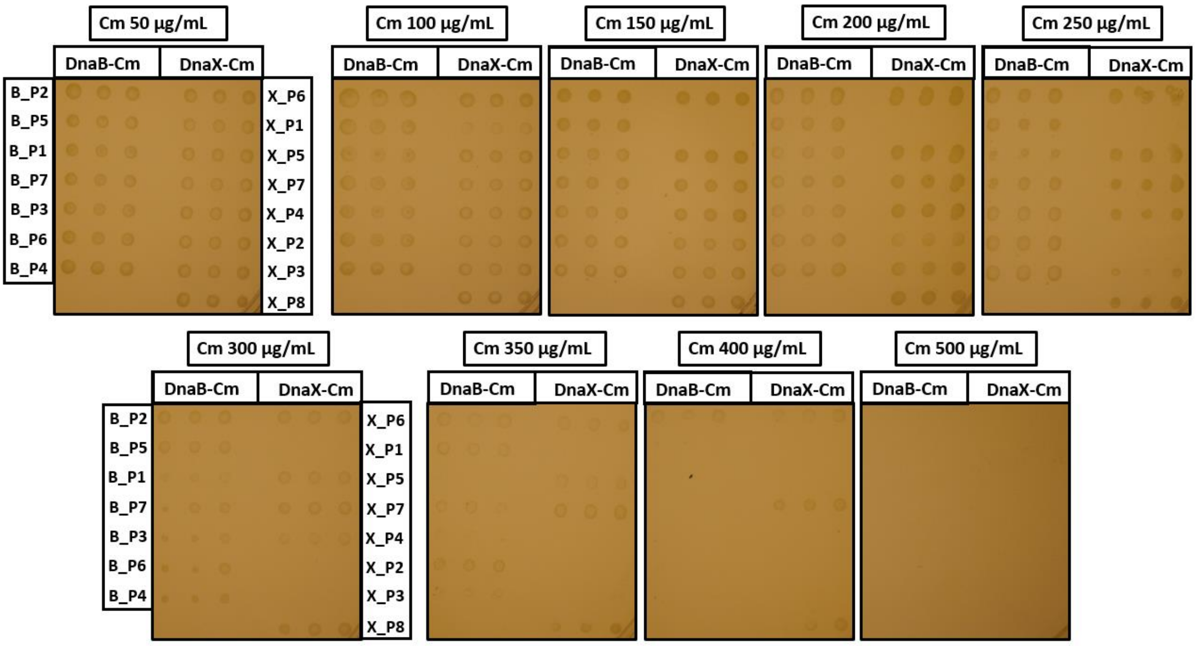

#### <u>Supplementary Figure S2: Chloramphenicol gradient for *DnaB-cat* and *DnaX-cat*</u>

E. coli strains carrying the *DnaB-cat* and *DnaX-cat* with the different GES were grown overnight in 96 well plates, diluted 100 times and replica plated on LB-agar plates containing increasing chloramphenicol concentrations. The plate concentration is indicated above each pictures. The GES of *DnaB-cat* and *DnaX-cat* are indicated on the side of the first image.

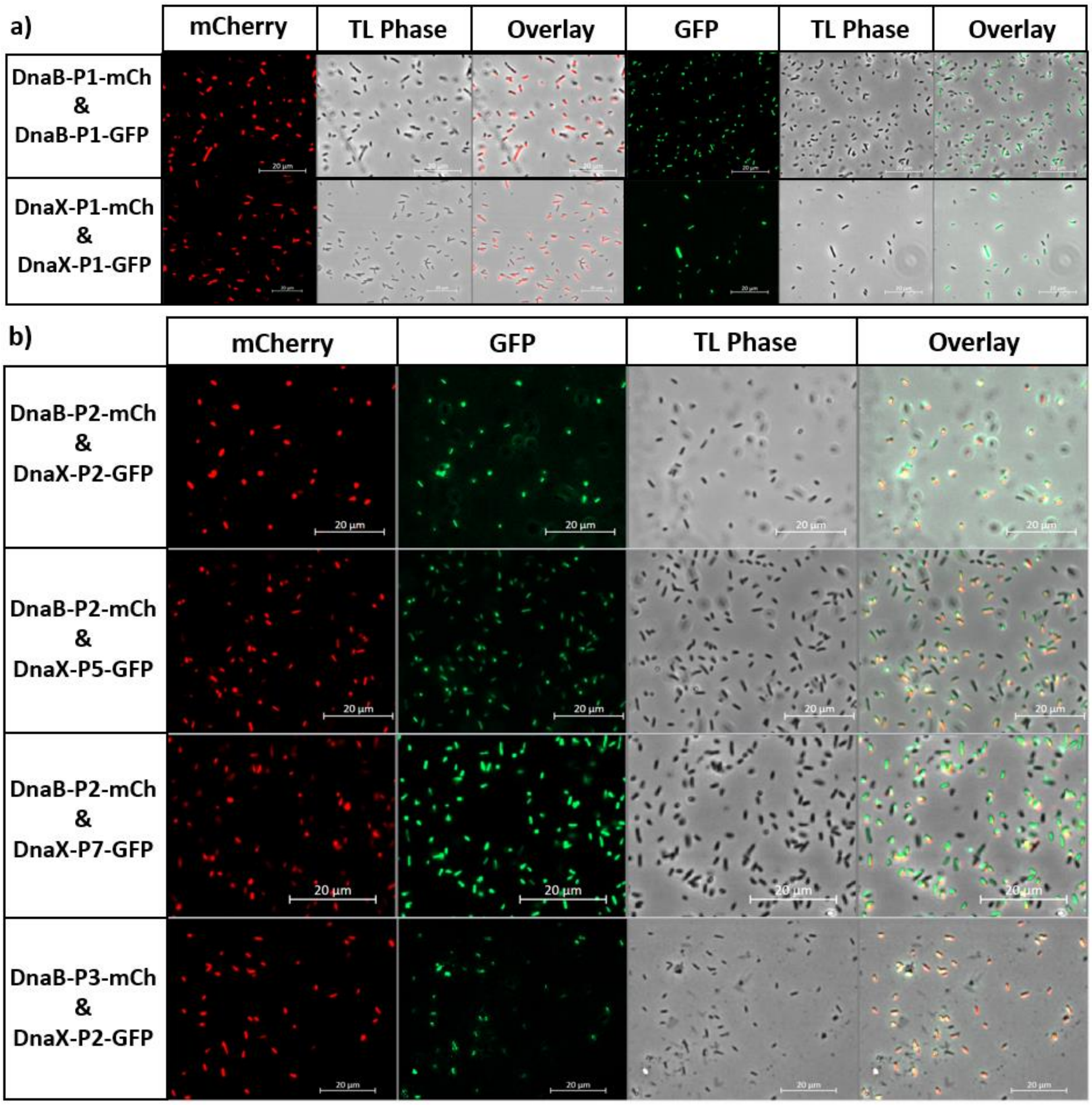

**Supplementary Figure S3: Fluorescence microscopy of weak GES and coupled SIGER systems**

a) Fluorescence microscopy of the weakest GES of *DnaB* and *DnaX* (B_P1 and X_P1 resp.) expressing GFP and mCherry separately.

b) The coupled SIGER systems were also analyzed to confirm the fluorescence of the weakest used GES (B_P2 and X_P2 resp.) and the co-localization of mCherry and GFP in *E. coli* cells. The fluorescence of mCherry, GFP, the TL phase and the overlay is presented for each constructs (see *Methods* for parameters).

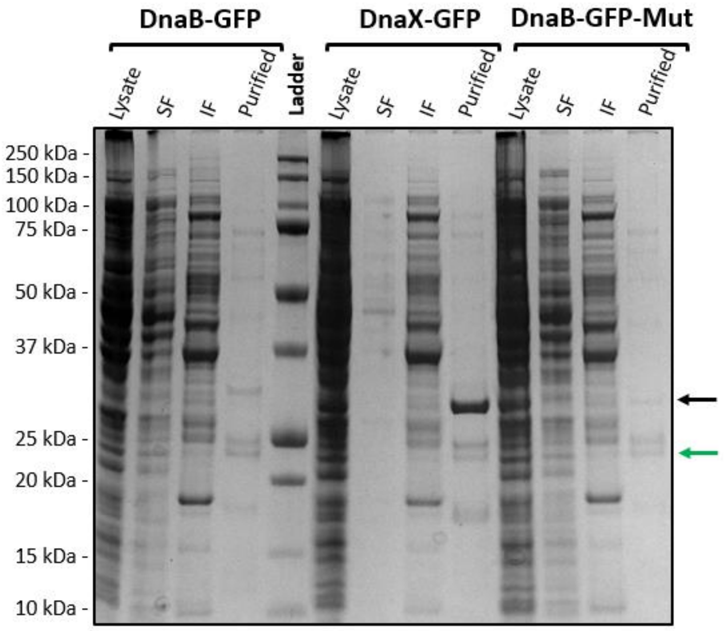

**Supplementary Figure S4: SDS-PAGE of purified proteins prior to proteomics analysis**

*E. coli* carrying P7_*DnaB-GFP*, P7_*DnaB_mut-GFP* and P7_*DnaX-GFP* were grown in flasks and their protein content was analyzed on SDS-PAGE. Cells were disrupted by sonication (lysate), cell debris (insoluble fraction IF) were removed and the soluble fraction (SF) was purified on Ni-Sepharose column. Each fraction was loaded on SDS-PAGE as indicated above the image. The black arrow is pointing at the bands corresponding to uncleaved Intein-GFP protein and the green arrow points at the released GFP protein. The bands corresponding to the uncleaved products and the GFP band were excised for proteomics analysis. The proteomics analysis of the bands confirmed the respective amino acid sequences.

| | |
|---|---|
| **DnaB-GFP** | MRESGAISGDSLISLASTGKRVSIKDLLDEKDFEIWAINEQTMKLESAKVSRVFCTGKKLVYILKT<br>RLGRTIKATANHRFLTIDGWKRLDELSLKEHIALPRKLESSSLQLSPEIEKLSQSDIYWDSIVSITET<br>GVEEVFDLTVPGPHNFVANDIIVHNSRGP<br>RKGEELFTGVVPILVELDGDVNGHKFSVRGEGEGDATNGKLTLKFICTTGKLPVPWPTLVTTLTY<br>GVQCFARYPDHMKQHDFFKSAMPEGYVQERTISFKDDGTYKTRAEVKFEGDTLVNRIELKGID<br>FKEDGNILGHKLEYNFNSHNVYITADKQKNGIKANFKIRHNVEDGSVQLADHYQQNTPIGDGP<br>VLLPDNHYLSTQSVLSKDPNEKRDHMVLLEFVTAAGITHGMDELYKHHHHHH* |
| **DnaB-mut-GFP** | MRESGAISGDSLISLASTGKRVSIKDLLDEKDFEIWAINEQTMKLESAKVSRVFCTGKKLVYILKT<br>RLGRTIKATANHRFLTIDGWKRLDELSLKEHIALPRKLESSSLQLSPEIEKLSQSDIYWDSIVSITET<br>GVEEVFDLTVPGP<br>RKGEELFTGVVPILVELDGDVNGHKFSVRGEGEGDATNGKLTLKFICTTGKLPVPWPTLVTTLTY<br>GVQCFARYPDHMKQHDFFKSAMPEGYVQERTISFKDDGTYKTRAEVKFEGDTLVNRIELKGID<br>FKEDGNILGHKLEYNFNSHNVYITADKQKNGIKANFKIRHNVEDGSVQLADHYQQNTPIGDGP<br>VLLPDNHYLSTQSVLSKDPNEKRDHMVLLEFVTAAGITHGMDELYKHHHHHH* |
| **DnaX-GFP** | MNGLMSIDNPQIKGREVLSYNETLQQWEYKKVLRWLDRGEKQTLSIKTKNSTVRCTANHLIRT<br>EQGWTRAENITPGMKILSPASASGHHHHHHGGSGSSPQWHTNFEEVESVTKGQVEKVYDLE<br>VEDNHNFVANGLLVHNSRGP<br>RKGEELFTGVVPILVELDGDVNGHKFSVRGEGEGDATNGKLTLKFICTTGKLPVPWPTLVTTLTY<br>GVQCFARYPDHMKQHDFFKSAMPEGYVQERTISFKDDGTYKTRAEVKFEGDTLVNRIELKGID<br>FKEDGNILGHKLEYNFNSHNVYITADKQKNGIKANFKIRHNVEDGSVQLADHYQQNTPIGDGP<br>VLLPDNHYLSTQSVLSKDPNEKRDHMVLLEFVTAAGITHGMDELYKHHHHHH* |
| **DnaX-mut-GFP** | MNGLMSIDNPQIKGREVLSYNETLQQWEYKKVLRWLDRGEKQTLSIKTKNSTVRCTANHLIRT<br>EQGWTRAENITPGMKILSPASASGHHHHHHGGSGSSPQWHTNFEEVESVTKGQVEKVYDLE<br>VEDNPT* |
| **DnaX-mut-GFP in-frame GFP** | MTWRWKIIP<br>RKGEELFTGVVPILVELDGDVNGHKFSVRGEGEGDATNGKLTLKFICTTGKLPVPWPTLVTTLTY<br>GVQCFARYPDHMKQHDFFKSAMPEGYVQERTISFKDDGTYKTRAEVKFEGDTLVNRIELKGID<br>FKEDGNILGHKLEYNFNSHNVYITADKQKNGIKANFKIRHNVEDGSVQLADHYQQNTPIGDGP<br>VLLPDNHYLSTQSVLSKDPNEKRDHMVLLEFVTAAGITHGMDELYKHHHHHH* |
| **GFP** | SRGPRKGEELFTGVVPILVELDGDVNGHKFSVRGEGEGDATNGKLTLKFICTTGKLPVPWPTLV<br>TTLTYGVQCFARYPDHMKQHDFFKSAMPEGYVQERTISFKDDGTYKTRAEVKFEGDTLVNRIE<br>LKGIDFKEDGNILGHKLEYNFNSHNVYITADKQKNGIKANFKIRHNVEDGSVQLADHYQQNTPI<br>GDGPVLLPDNHYLSTQSVLSKDPNEKRDHMVLLEFVTAAGITHGMDELYKHHHHHH* |

**Supplementary Figure S5: Proteomics sequence coverage of analyzed bands from SDS-PAGE**

The table presents the amino acid sequences of the proteins resulting from the expression of *DnaB-GFP*, *DnaX-GFP* and their mutated versions. The N-terminal asparagine responsible for cyclization and autocatalytic release of the POI is underline in blue. The linker peptide that connects the inteins and the POIs in highlighted in gray. The sequence of GFP is highlighted in green. Bands extracted from the SDS-PAGE were analyzed by proteomics. Peptides found in the proteomics analysis are represented with red letters. Alternative translation initiation start sites are highlighted in red.

The 45- and 43-kDa proteins corresponding to the uncleaved DnaB-GFP and DnaX-GFP proteins were confirmed by the proteomics analysis. The 28-kDa protein was confirmed to be the released GFP protein. The DnaB_mut-GFP analysis demonstrated the presence of an alternative translation initiation start site possibly at V136. The DnaX_mut-GFP mRNA also presents an in-frame methionine nine amino acids before the start of GFP that produces an unexpected N-terminal elongated GFP.

**Supplementary Table S4. Overview of protein yields and cleavage percentages for the three tested proteins.** Cleavage percentages are estimated from the band intensities on SDS-PAGE gel.

| Protein | Growth temperature | Total protein yield; cleaved and uncleaved (mg/L of culture) | Cleavage (%) |
|---|---|---|---|
| GFP | 22°C | 5.0 | 43 |
| | 37°C | 32.1 | 64 |
| GumM | 22°C | 3.6 | 51 |
| CBM73 | 22°C | 2.9 | 37 |

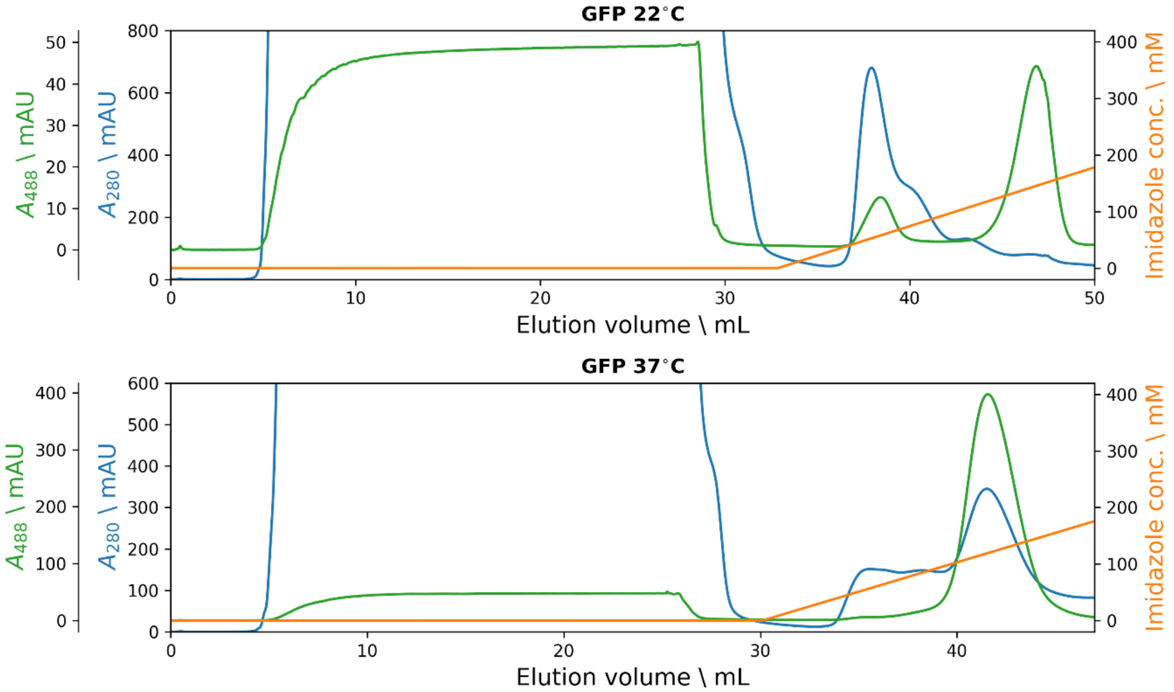

**Supplementary Figure S6. Purification of GFP from cultures grown at 22 °C and 37 °C.** The figure shows chromatograms from the purifications carried out using a 1 mL HisTrap HP Ni-sepharose column. The blue curves show absorbance at 280 nm (A280), the green curves show absorbance at 488 nm (A488), and the orange curves show the gradient concentration of imidazole used during elution. Flow-through (unbound protein) is found between 0 – 30 mL, whereas elution starts after 30 mL. For illustration purposes, the flow-through peak (reaching a maximum of 3000 mAU) is truncated. Note that the scale of the absorbance axes is different for the two temperatures, showing a significantly larger GFP yield at 37°C.
